# Modelling human social vision with cinematic stimuli

**DOI:** 10.1101/2024.10.18.618846

**Authors:** Severi Santavirta, Birgitta Paranko, Kerttu Seppälä, Jukka Hyönä, Lauri Nummenmaa

## Abstract

Sociability is central for humans. Visual information ranging from low-level physical features (e.g. luminance) to semantic information (e.g. face recognition) and high-level social inference (e.g. emotional valence of social interactions) is constantly sampled for navigating the social world. Here we utilize large-scale eye tracking during natural vision for mapping how different levels of visual information guide the perception of social scenes. In three experiments, participants (N = 166) watched full-length films and short movie clips with varying social content (total duration: 193 minutes) during eye tracking. To model the association between the perceptual features and spatiotemporal eye movement parameters (gaze position, gaze synchronization, pupil size and blinking), we extracted 39 stimulus features from the movies including low-level audiovisual features (e.g. luminance, motion), presence and location of mid-level semantic categories (e.g. faces, objects) and high-level social information (e.g. body movements, pleasantness). Pupil size was modulated by luminance, scene cuts and emotional arousal while gaze position was most accurately predicted by a combination of the presence of human faces, local motion and entropy. Faces and eyes were prioritized over other semantic categories and blinking rate decreased during periods of attentional engagement. Altogether the results show that human social vision is primarily guided by low-level physical features and mid-level semantic categories, while high-level social features such as emotional arousal primarily modulate pupillary responses.

## Introduction

Visual information about other people is central for humans. The attentional systems constantly adjust the sampling of visual information to extract features that have the highest processing priority at each moment. This is primarily achieved by controlling the gaze position, and also via modulating visual sampling by adjusting fixation frequency, segmenting the visual input into chunks through blinking, as well as by controlling for the number of photons landing on the retina through pupillary control. This cascade of operations ensures that we, for example, quickly recognize affective content from scenes (Nummenmaa et al., 2010) and facial expressions (Calvo & Nummenmaa, 2008) and preferentially allocate attention to social signals ranging from faces to reproductive cues (Morrisey et al., 2019).

Modelling of human social vision is complicated by the complex spatiotemporal dynamics of gaze behaviour. Mapping the links between high-dimensional stimulus models during natural vision is difficult for traditional eye-tracking paradigms because isolating the contribution of single social features (e.g. emotional content) to gaze control is complicated by the overlapping and hierarchical time scales, parallel processing of sensory features, and ever-changing locations of the objects of interest in dynamic scenes. This often necessitates analyzing of spatial gaze data during artificial static snapshots of scenes in pictures, which however precludes the analysis of the effects of temporally varying stimulus properties on gaze control. It has thus been debated whether the results from such simplified conditions transfer to social vision operating in the dynamic social environment (Williams & Castelhano, 2019), particularly as the visual system is differently influenced by static versus dynamic stimuli (Dorr et al., 2010). Furthermore, different external stimulus features may have unique, additive or interactive effects on spatial and temporal eye movement parameters (e.g. pupillary response, gaze position, blinking, etc.); yet, these features are rarely analyzed in unitary models. Instead, studies have mainly incorporated simple designs investigating one or few components of the visual system (e.g. pupil size or gaze control) for one or few phenomena at a time. Currently, our understanding of moment-to-moment high-dimensional modelling of the simultaneous impact of different external features on dynamic scenes is lacking. This kind of dynamic mapping of the links between high-dimensional social feature space and different eye movement metrics would require a holistic and data-driven approach which we present in the current research.

Social information processing is organized along eight main perceptual dimensions, such as the perception of pleasant versus unpleasant situations, social hierarchy, or communication (Santavirta et al., 2024), and functional neuroimaging has established that large-scale occipito-temporal brain networks are involved in processing of these features (Lahnakoski et al., 2012; Santavirta et al., 2023). It however remains unresolved how these social features are extracted through the visual attentional systems. The human visual system could be merely tuned to follow regularities in simple physical properties such as low-level visual features (e.g. luminance) or mid-level semantic categorization (e.g. faces), and complex social inference is a mere result derived from these inputs. Alternatively, high-level social information could be used in guiding social vision in top-down manner during dynamic perception along with the low- and mid-level perceptual features. Studying simultaneous effects of low-, mid-, and high-level stimulus features on the visual system would be required to distinguish these alternative possibilities.

Gaze is the most studied component of the visual system. During dynamic vision, the gaze positions synchronize between individuals (Dorr et al., 2010; Franchak et al., 2016; Smith & Mital, 2013). The synchronization of gaze indicates shared attention which is useful for common understanding of the events and for cooperation. Synchronization can be measured using intersubject correlation (ISC), which can be as high as 0.4 – 0.6 during movie watching (Hasson et al., 2008; Wang et al., 2012) indicating high attentional coherence and a major influence of the stimulus properties in directing the gaze. A large bulk of studies has addressed how gaze is directed by external or intrinsic features during scene perception, and both bottom-up and top-down models for gaze control have been proposed (J. M. Henderson, 2003). Bottom-up saliency models predict gaze direction reasonably well based on saliency maps computed from color, intensity, and orientation information in images (Itti & Koch, 2000). However, these pure bottom-up models cannot explain innate strong human preferences for specific semantic object categories such as human faces and eyes that are not perceptually particularly salient (E. Birmingham et al., 2009; Elina Birmingham et al., 2008). Debate whether gaze directions are best modelled with low-level saliency or object-based models is ongoing (Stoll et al., 2015), and the latest evidence suggests that saliency and object-information together yield better predictive models than either of them alone (Nuthmann et al., 2020; Roth et al., 2023). Recent deep learning models for gaze patterns can yield high predictive performance without predefined features for images (Cornia et al., 2018; Lou et al., 2022) and some also for videos (Bellitto et al., 2021; Jain et al., 2021). These complex models do not distinguish between bottom-up and top-down influences, since complex models can learn to detect specific objects (for example faces) from local spatial features. However, it is difficult to know exactly what information these models have learned to use when predicting scene saliency. Indirect methods can be utilized to interpret the outputs of these complex models (Hayes & Henderson, 2021), but currently they offer little theoretical insight about the visual system in operation.

Gaze direction is not the only component of the visual system indexing perceptual and cognitive processing. Pupil automatically regulates the flow of light on the retina, and the pupil size also fluctuates as a function of internal states such as emotional arousal and cognitive effort. Seminal results indicated that pupil dilates when watching pleasant images (Hess & Polt, 1960), while recent studies have shown that pleasant and unpleasant emotions trigger pupil dilation when viewing images (Bradley et al., 2008), listening to sounds (Oliva & Anikin, 2018; Partala & Surakka, 2003), and also when imagining emotional episodes (R. R. Henderson et al., 2018). This arousal-driven pupillary response is usually stronger for unpleasant than pleasant stimuli (Babiker et al., 2013; Kawai et al., 2013). Pupil dilation is also associated with cognitive effort in various cognitive control tasks (Ayres et al., 2021; Hyönä et al., 1995; Kahneman & Beatty, 1966; van der Wel & van Steenbergen, 2018). In turn, pupil has been found to constrict as a function of attractiveness of faces and natural scenes (Liao et al., 2021). Pupil constriction predicts how well people memorize images and reveals the novelty of a scene (Naber et al., 2013). Pupil seems to constrict even during sudden transitions of simple stimuli in isoluminous conditions (Kimura et al., 2014). These luminance-independent pupillary responses are likely mediated by the adrenergic and cholinergic neurotransmitter systems that are engaged during both emotions and cognitive effort to maximize attentional prioritization and arousal levels in response to environmental demands (Joshi et al., 2016; Reimer et al., 2016). Real-life pupillary response is a complex combination of pupillary light reflex and higher-order effects such as those induced by emotions (Cherng et al., 2020; Steinhauer et al., 2004), but it is not yet established how multiple factors simultaneously modulate the pupil size during natural vision.

Blinking is another component of the visual system that can reveal cognitive processes outside of its main function. Blinks keep the eyes lubricated and remove irritants from the eye surface but the automatic eyeblink reflex is also a part of the startle response, where the orbicularis oculi muscle contracts rapidly after brief and intense auditory, visual, or tactile stimuli (Grillon & Baas, 2003). Blinking is however also influenced by cognitive and affective factors. Blinking and pupil dilation are suggested to be the most sensitive physiological measures of cognitive load (Ayres et al., 2021). In a naturalistic study on *Mastermind* TV-quiz contestants, blinking was strongly modulated by the task so that more blinks occurred at attentional or cognitive breakpoints right before and right after each question, contestant’s response or feedback (Wyly et al., 2024). In addition, blink rate varies as a function of the emotional content indicating motivational relevance and biological significance of the stimuli (Maffei & Angrilli, 2019). Blinking is found to synchronize between participants during shared natural vision indicating shared attention (Nakano et al., 2009); moreover, blink synchronization is higher among participants who are more interested in the stimuli (Nakano & Miyazaki, 2019). Thus, blinking reflects cognitive load, attention, or interest levels of the participants, and possibly the emotional context. Functional neuroimaging results have also established a link between blinking and attentional engagement, as decreased brain activity in the dorsal attention network after blink onset has been identified simultaneously with increased activity in the default-mode network (Nakano et al., 2013).

### The current study

To understand how social vision is controlled during natural perception, a unified approach linking spatial and temporal aspects of eye movements to low- and high-level scene features is needed. The objective of this study was to establish a unified framework for how human visual system is tuned by external features in dynamic social scenes. We studied moment-to-moment gaze positions, pupillary response, and blinking behaviour while participants freely viewed social movies. We aimed to simultaneously investigate the contributions of low-level physical features, semantic perceptual features, and high-level perceived social information on the spatial and temporal dynamics of human social vision. Movie scenes were utilized as dynamic stimuli since they depict highly engaging social and emotional contents and effectively synchronize human gaze positions (Hasson et al., 2008), which makes them ideal naturalistic stimuli used in the laboratory (Adolphs et al., 2016; Saarimäki, 2021).

We conducted three experiments with a total of 166 participants and 193 minutes of movie stimuli to ensure sufficient statistical power and to allow replicability testing for results across the datasets. A total of 39 different perceptual features from low-level audiovisual properties (e.g. luminance, motion) to mid-level perceptual semantic objects (e.g. eyes, objects) and high-level perceived socioemotional features (e.g. pleasantness, talking) were dynamically extracted from the movies. A stimulus model including 16 predictors was generated from the extracted stimulus features after studying the covariance structure of the original 39 features. Four main analysis techniques (total gaze time analysis, multi-step regression analysis, gaze prediction analysis, and scene cut effect analysis) were developed to link the extracted stimulus features with the eye-tracking parameters.

Our results show that human social vision is primarily guided by low-level physical features and mid-level semantic categories, while high-level social features, such as emotional arousal, primarily modulate pupillary responses. Importantly, gaze direction, pupillary responses and blinking all indexed unique response patterns in relation to external stimulus features in dynamic perception.

**Table 1.** Demographic information of the participant sample.

| Experiment | Subjects | M/F | Mean age (range) |
| --- | --- | --- | --- |
| Experiment 1, Movie clips | 106 | 40/66 | 27.1 (19 - 74) |
| Experiment 2, Feature film | 21 | 2/19 | 23.6 (19 - 38) |
| Experiment 3, Horror movie | 24 | 5/19 | 27.0 (19 - 57) |

## Results

### Eye movements during movie viewing

Figure 1 shows the histograms for fixations, saccades, and blinks during movie viewing. The data are combined over all three experiments. The fixation, saccade and blink durations were similar across the experiments (**Figure SI-1**). The median fixation duration was 309 ms (q5% - q95%: 135 ms – 1047 ms), the median saccade duration was 27 ms (q5% - q95%: 11 ms – 55 ms) and the median blink duration was 91 ms (q5% - q95%: 23 ms – 217 ms). The fixation, saccade and blink rates varied between participants. The median fixation rate was 2.12 / s (q5% - q95%: 1.77 / s – 2.52 / s) and the median blink rate was 0.20 / s (q5% - q95%: 0.05 / s – 0.55 / s). The median total fixation time (total duration of fixations / total stimulus duration) was 0.89 (q5% - q95%: 0.82 – 0.93) and the median total blink time (total duration of blinks / total stimulus duration) was 0.02 (q5% - q95%: 0.00 – 0.07). The gaze patterns were moderately consistent across subjects (Mean ISC: 0.37, SD: 0.20). The eye-tracking measures (pupil size, ISC, fixation rate and blink synchronization) were mostly uncorrelated with each other (Figure 2 **& Figure SI-2**). In the 200 and 500 ms temporal scale, a negative correlation was observed between fixation rate and ISC (r = -0.14) and between blinking and ISC (r = -0.09) consistently across the three experiments, but this weak correlation was not consistent across datasets when analyzed in longer temporal scales (1000 ms – 4000 ms).

**Figure 1.**
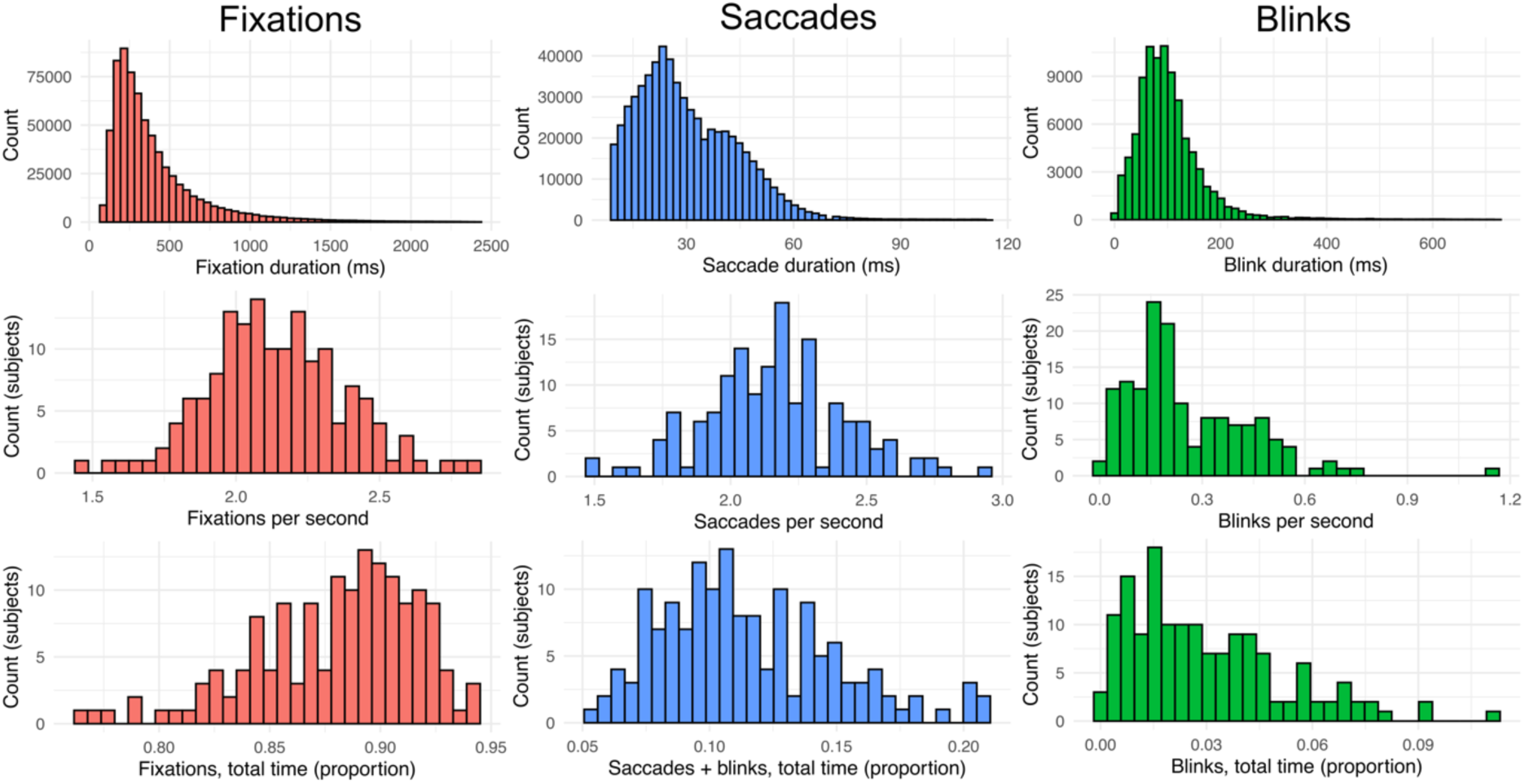
Distributions of fixations, saccades, and blinks during movie viewing. The top row shows distributions for fixation, saccade and blink durations over all participants and datasets. The middle row shows the distributions of participant-wise mean fixation / saccade / blink rates and the bottom row shows the distributions of participant-wise proportions of time allocated for fixations, saccades, or blinks.

**Figure 2.**
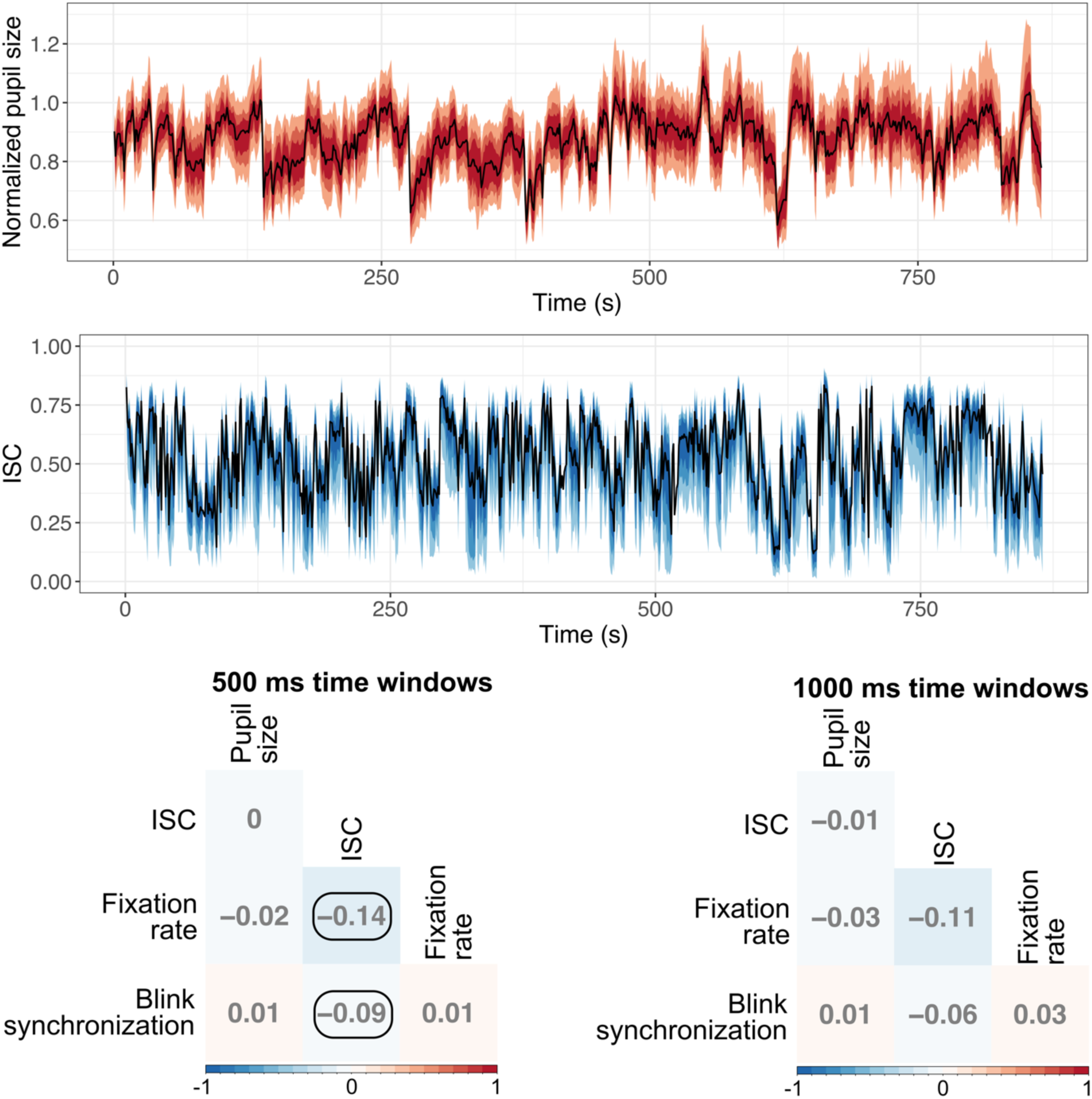
Top: Time series of median pupil size and ISC across subjects in the Experiment 1 data (coloured areas mark the 40%, 60%, and 80% quantile intervals, see **Figure SI-3** for time series of Experiment 2 & 3 data). Bottom: correlations between the eye-tracking parameters. The correlations were calculated for each experiment separately and then averaged over datasets. Correlations that had consistent signs over all experiments are circled. The correlations were calculated in multiple different time scales (200 ms – 4000 ms, see **Figure SI-2**). Blink synchronization indicates how many participants blinked in a given time window.

### Gaze time analysis

The participants showed an attentional preference in the dynamic stimuli to the eye and mouth area in all three datasets (Figure 3). The eyes were viewed 21% - 33% of the total stimulus time, which was 16 – 24 percentage points more than would be expected by chance (p < 0.005). The mouth area was viewed 10% - 11% of the time, which was 7 – 8 percentage points more than would be expected by chance (p < 0.005). The bodies were looked at 18% - 28% of the total time, which was 4 – 7 percentage points less than would be expected by chance (p < 0.005). Similarly, background was viewed 15% - 23% of the total time, which was 17 – 23 percentage points less than would be expected by chance (p < 0.005). The difference in gaze time for objects and faces excluding the eye and mouth area was small, and the direction of difference was inconsistent between the datasets. People also gazed at unclassified areas, for which the computer vision algorithm failed to assign a reliable category for 10% - 14% of the time which, was 1% - 2% less than would be expected by chance (p < 0.005). This indicates that the computer vision algorithm did not miss highly prioritized information from the stimuli. People gazed at areas outside the video less than 1% of the time. Since the real gaze coordinates were randomly sampled for the permuted chance time estimation, the gaze time outside the video area cannot be differentiated from chance with the implemented test.

**Figure 3.**
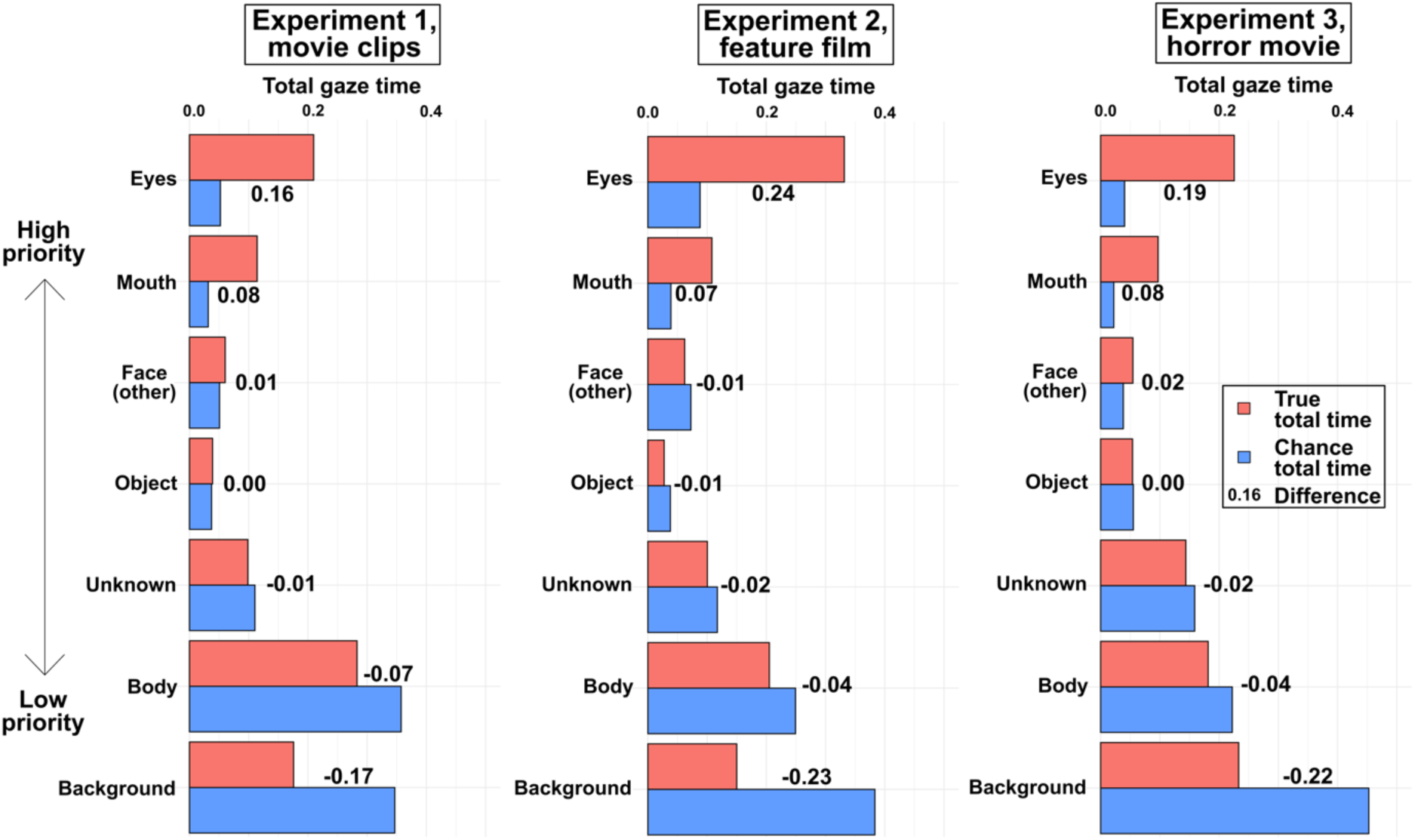
Results of the gaze time analysis. The gaze time analysis results are plotted separately for each experiment. The red bars show the true total gaze times (proportional to the experiment duration), while the blue bars show the total times that would be expected by chance for different semantic classes. The numbers show the difference between true gaze time and estimated chance gaze time. All differences between true and chance gaze times were statistically significant based on a permutation test (p < 0.005).

### Multi-step regression analysis

We next established links between spatiotemporal gaze parameters and low-, mid-, and high-level stimulus features. To that end, pupil size, ISC, fixation rate and blink rate were modeled with a stimulus model of 16 low-level, semantic, and social perceptual features using a multi-step regression approach. The stimulus model was generated from the extracted 39 features based on the covariance structure between the original features. Figure 4 summarizes the regression results for the representative 500 ms temporal analysis scale. The results were, however, consistent when the analyses were repeated across different time scales (**Figure SI-4**). While most of the predictors had a consistent sign of association between the eye-tracking variables in simple regression over the cross-validation rounds (leave-one-experiment-out cross-validation, not tested for statistical significance), only a limited number of predictors increased the subsequent multiple regression models’ out-of-sample prediction accuracies significantly.

**Figure 4.**
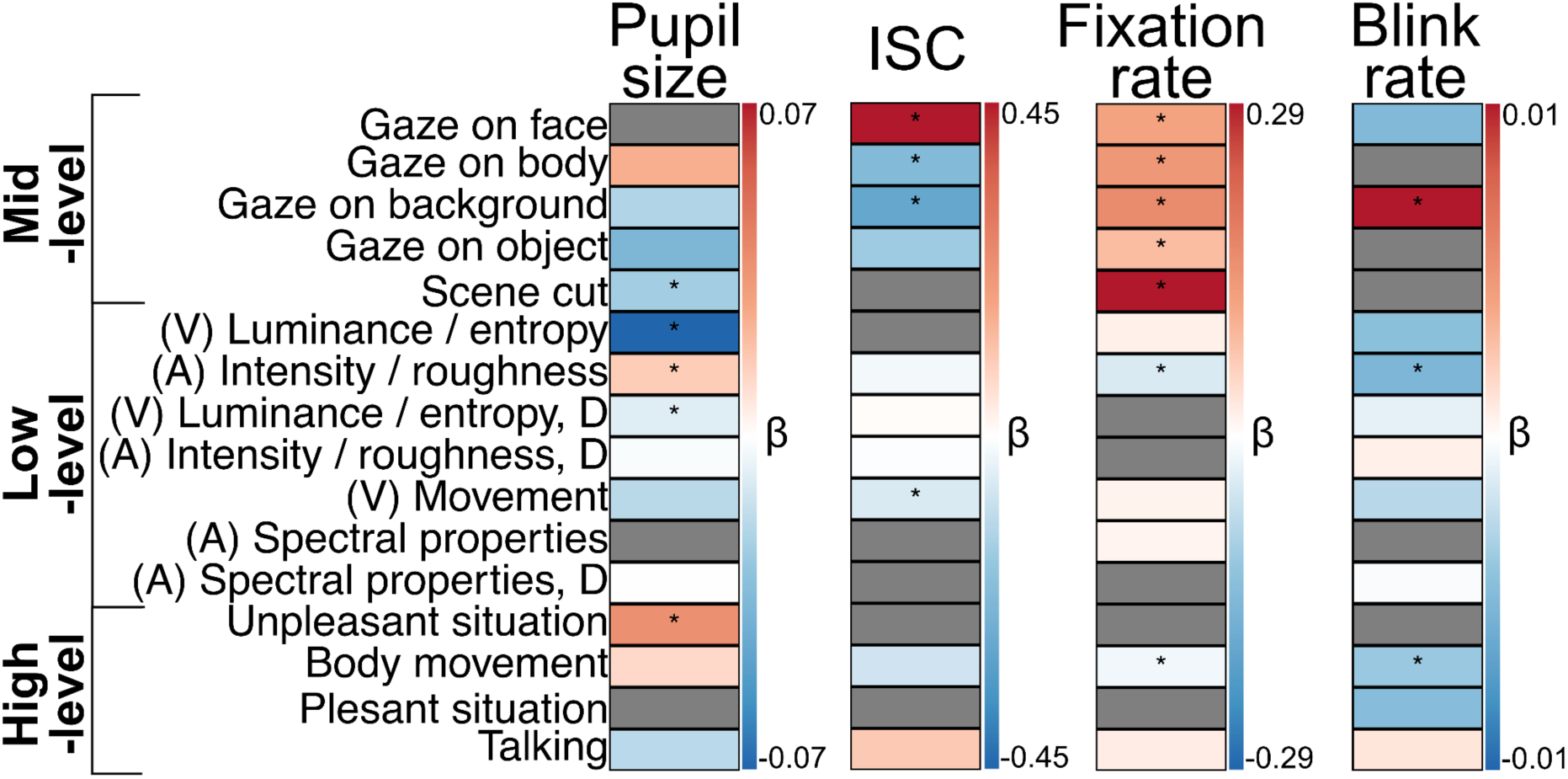
The results of multi-step regression analyses for pupil size, ISC, fixation rate and blink rate in the 500 ms time scale. Coefficients from the simple regressions are visualized with a blue (negative) – red (positive) colour gradient. Grey colour indicates that the direction of the association was inconsistent between the simple regression cross-validation rounds (leave-one-experiment-out cross-validation). The final multiple regression model included the features marked with asterisks. These features increased the model’s out-of-sample prediction accuracy significantly compared to chance (p < 0.05) when added to a multiple regression model. (V) denotes low-level visual predictors, (A) denotes auditory low-level predictors and D denotes the time derivative of the feature.

**Pupil size** was most consistently associated with low-level features and perceived unpleasantness (Figure 4 **& Figure SI-4**). In the 500 ms time scale, adding consistent predictors one-by-one into a multiple regression model starting from the predictor with the best out-of-sample prediction accuracy in simple regression, the predictors “Luminance / entropy” (negative association), “Luminance / entropy, time derivative” (negative association), “Audio intensity / roughness” (positive association), “Scene cut” (negative association) and “Unpleasant situation” (positive association) increased the model’s out-of-sample prediction accuracy significantly compared to the permuted chance level (p < 0.05). The final models including these five features produced predictions in the test set (leave-one-experiment-out) that correlated highly with the true pupil size (Exp. 1 as the test set: r = 0.36, Exp. 2 as the test set: r = 0.49, Exp. 3 as the test set: r = 0.50).

**ISC** of gaze was most consistently associated with mid-level features and motion (Figure 4 **& Figure SI-4**). In the 500 ms time scale, the predictors “Gaze on face” (positive association), “Gaze on body” (negative association), “Gaze on background” (negative association) and “Visual movement” (negative association) increased the multiple regression model’s out-of-sample prediction accuracy significantly (p < 0.05). The final models including these four features produced predictions in the test set (leave-one-experiment-out) that correlated highly with the true ISC (Exp. 1 as the test set: r = 0.43, Exp. 2 as the test set: r = 0.33, Exp. 3 as the test set: r = 0.40).

**Fixation rate** was most consistently associated with mid-level features and audio intensity / roughness (Figure 4 **& Figure SI-4**). In the 500 ms time scale, the predictors “Scene cut” (positive association), “Gaze on face” (positive association), “Gaze on body” (positive association), “Gaze on background” (positive association), “Gaze on object” (positive association), “Audio intensity / roughness” (negative association) and “Body movement” (negative association) increased the multiple regression model’s out-of-sample prediction accuracy significantly (p < 0.05). The final models including these seven features produced predictions in the test set (leave-one-experiment-out) that correlated moderately with the true fixation rate (Exp. 1 as the test set: r = 0.26, Exp. 2 as the test set: r = 0.21, Exp. 3 as the test set: r = 0.24).

**Blink rate** was mainly associated with “Gaze on background” (positive association), “Audio intensity / roughness” (negative association) and “Body movement” (negative association). These variables increased the model’s out-of-sample prediction accuracy significantly (p < 0.05) in the 500 ms time scale. The final models including these three features were not able to predict much variation in the leave-one-experiment-out test set (Exp. 1 as the test set: r = 0.03, Exp. 2 as the test set: r = 0.02, Exp. 3 as the test set: r = 0.03).

All the above reported associations between perceptual features and eye-tracking measures were consistent between the initial simple regressions and the multi-step regression rounds indicating that the addition of other predictors in the model did not yield interpretational difficulties.

### Performance of the gaze prediction model

Random forest models using all available low-level, semantic, and social perceptual information were trained separately for each of the three datasets to predict the moment-to-moment population level gaze heatmaps. The performance of the trained models was tested with the two datasets that were left out from the model training. The performance of the models was evaluated by calculating the correlation between the true and predicted gaze heatmaps and by measuring how far, on average, the predicted peak value was from the true heatmap peak value (Figure 5). The evaluation metrics were consistent across the models trained with different datasets. Correlations between predicted heatmaps and true heatmaps in the testing dataset ranged between 0.41 and 0.47, while the peak value distances ranged between 10 % and 16 % of the image width indicating robust out-of-sample prediction performance.

**Figure 5.**
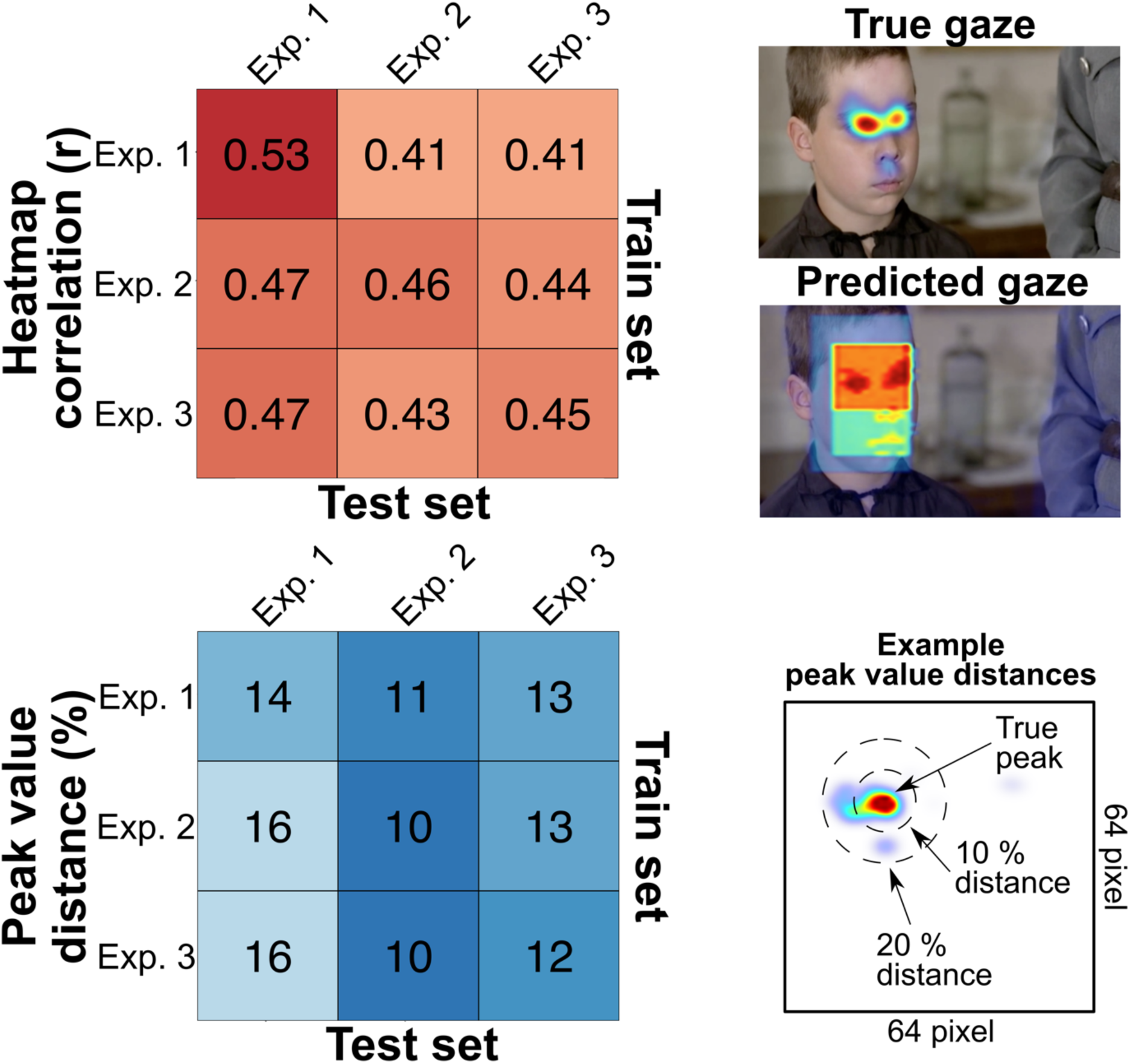
Performance of the gaze prediction model. The random forest model was trained with each dataset separately and the model’s performance was tested with the two other datasets. Top left: A confusion matrix showing the correlation of the true versus predicted gaze heatmaps for the trained models. Top right: True and predicted gaze heatmaps for one representative 200 ms time window. Bottom left: A confusion matrix showing how far, on average, the predicted peak heatmap value was from the true heatmap peak. Bottom right: Example peak value distances for 10 % and 20 % levels are visually provided for guiding the evaluation of the reported peak value distances. The sample images are from the Experiment 2 movie stimuli (Louhimies, 2008).

Relative importance measures how influential (between 0 and 1) a given predictor is for the model’s predictions. The relative importances were consistent across the models trained on different datasets (Figure 6, bar plot). The presence of eyes in a given voxel showed by far the highest importance for the predictions (Exp. 1: 0.48, Exp. 2: 0.67, Exp. 3: 0.46). After eyes, the presence of mouth (Exp. 1: 0.20, Exp. 2: 0.08, Exp. 3: 0.18), visual movement (Exp. 1: 0.09, Exp. 2: 0.10, Exp. 3: 0.06) and luminance / entropy (Exp. 1: 0.03, Exp. 2: 0.06, Exp. 3: 0.09) showed higher relative importances than the rest of the predictors.

**Figure 6.**
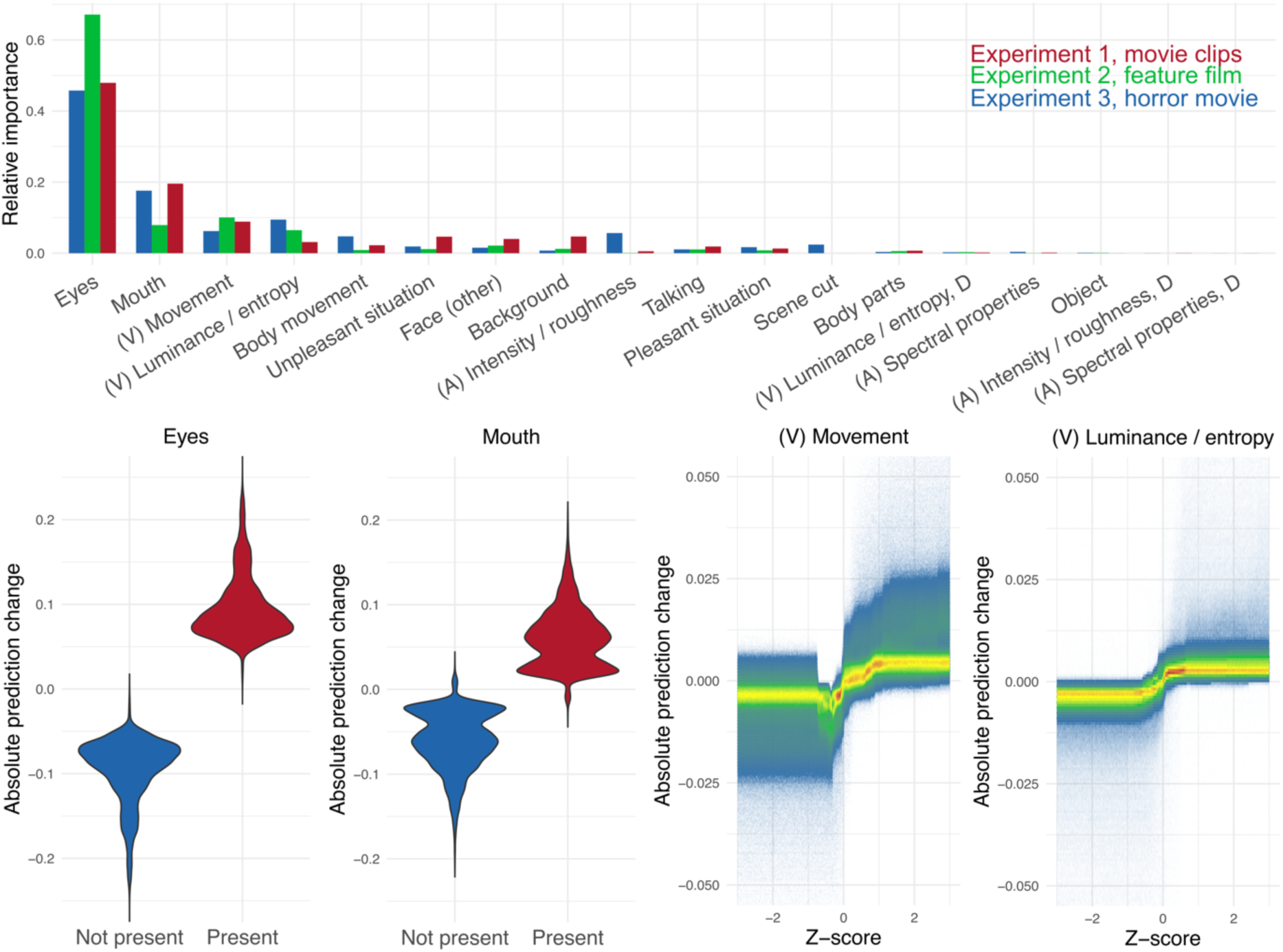
Interpretation of the gaze prediction models. Barplots visualize the relative importance of each predictor in the trained models. Violin plots for categorical and density plots for continuous predictors visualize the simulated influence of each predictor on the gaze probability prediction when other predictors’ values are held constant. (V) denotes low-level visual predictors, (A) denotes auditory low-level predictors and D denotes the time derivative of the feature.

To establish whether the important predictors have a positive or negative effect on the models’ predictions we simulated how the models’ predictions would change when all other predictors are held constant but the values of only one predictor are changed (Figure 6, violin and density plots). The random forest models predicted higher gaze probability when eyes or mouth were present than when they were absent. Additionally, higher than average local movement and visual luminance / entropy resulted in increased gaze probability prediction compared to lower-than-average values. The relative importance and change in simulated predictions with different predictor values was negligible for the rest of the predictors. The high-level social features are constant within each time window and cannot thus have an independent main effect on the gaze location. However, they could inform the model about the expected distributions of the gaze probabilities between time windows and high-level information can have an interaction effect with some pixelwise predictor (e.g. eyes are watched more closely in unpleasant situations). However, exploratory simulations of interactions between the perceived social predictors and the four most important predictors did not indicate any clear interaction effects.

### Scene change effect

Figure 7 shows the scene change effect on pupil size, ISC and blink synchronization. The findings were consistent across the datasets. Pupil size started to decrease rapidly after the scene changed and the pupil size was significantly decreased compared to the permuted baseline between 350 ms - 1150 ms after scene cut in all datasets (p < 0.05, based on a permutation test). The minimum pupil size (97 – 98 % of the baseline on average) was reached between 500 ms – 800 ms after the scene cut. ISC increased briefly after the scene changed. For continuous movies (Exp. 2 & 3), ISC increased significantly up to 800 ms after the scene transition (Exp. 2: between 400 ms and 800 ms after cut, Exp. 3: between 200 ms and 800 ms after cut, p < 0.05), while for short movie clips (Expr. 1) ISC remained elevated for a longer period of time (between 400 ms and 1400 ms after the scene cut, p < 0.05).

**Figure 7.**
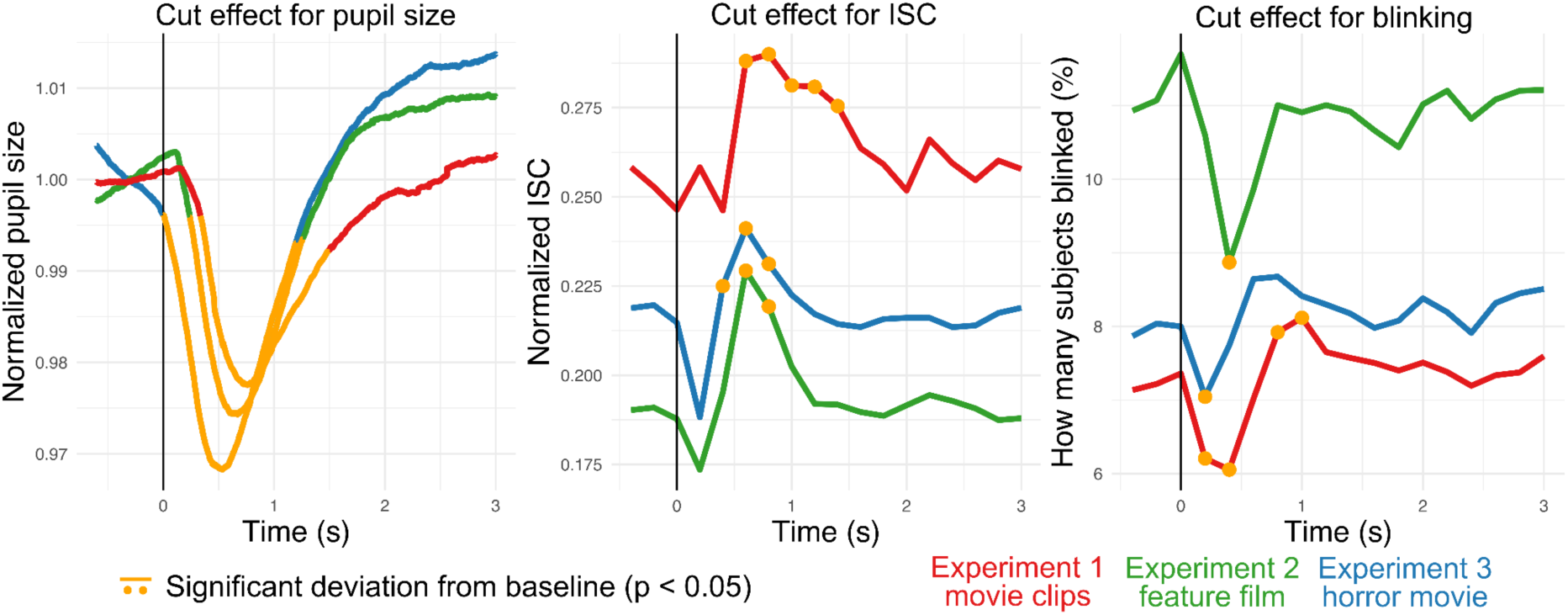
Temporal dynamics of pupil size, ISC and blinking after a scene cut separately for each experiment. The scene cut time is marked as a vertical line and the permuted statistically significant deviation period (pupil) or time points (ISC & blinking) from baseline are marked with orange (p < 0.05). Normalized ISC (ISC * stimulus area / total monitor area) is plotted since it account for the stimulus size differences between the Experiments (stimulus area / monitor area, Exp. 1: 0.53, Exp. 2: 0.94, Exp. 3: 0.95).

The percentage of participants that blinked in 200 ms time windows decreased shortly following the scene change in all datasets. For continuous movies, the percentage of participants blinking was briefly decreased at single time windows (Exp. 2: 200 ms – 400 ms after scene cut, Exp. 3: 0 ms – 200 ms after scene cut, p < 0.05). For the movie clip dataset (Exp. 1) blinking first decreased between 0 ms – 400 ms after the scene cut and then increased between 600 ms – 1000 ms (p < 0.05). The decrease followed by increase in blinking was not reliably observed in the movie datasets, although a similar increasing pattern after initial decrease in blinking was also suggested by the Experiment 3 data (Figure 7).

## Discussion

Our results established consistent relationships between low-level and semantic scene features on the spatiotemporal aspects of eye movements. During natural vision the gaze positions were relatively consistent across observers (mean ISC = 0.37) and mostly guided by mid-level semantic social information and low-level audiovisual features, whereas high-level social context contributed only minimally. In turn, pupil size was modulated by low-level sensory features as well as emotional content and scene transitions, while blinking rate was indicative of attentional enagement on the scenes. These results generalized across three independent datasets with different dynamic stimuli and participants. Our results thus suggest that gaze position, pupillary response and blinking are determined by independent, yet partially overlapping sets of external features, and that human social vision is primarily guided by low-level physical features and mid-level socially relevant semantic categories (most notably faces), while high-level socioemotional features such as emotional arousal primarily modulate pupillary responses (Figure 8).

**Figure 8.**
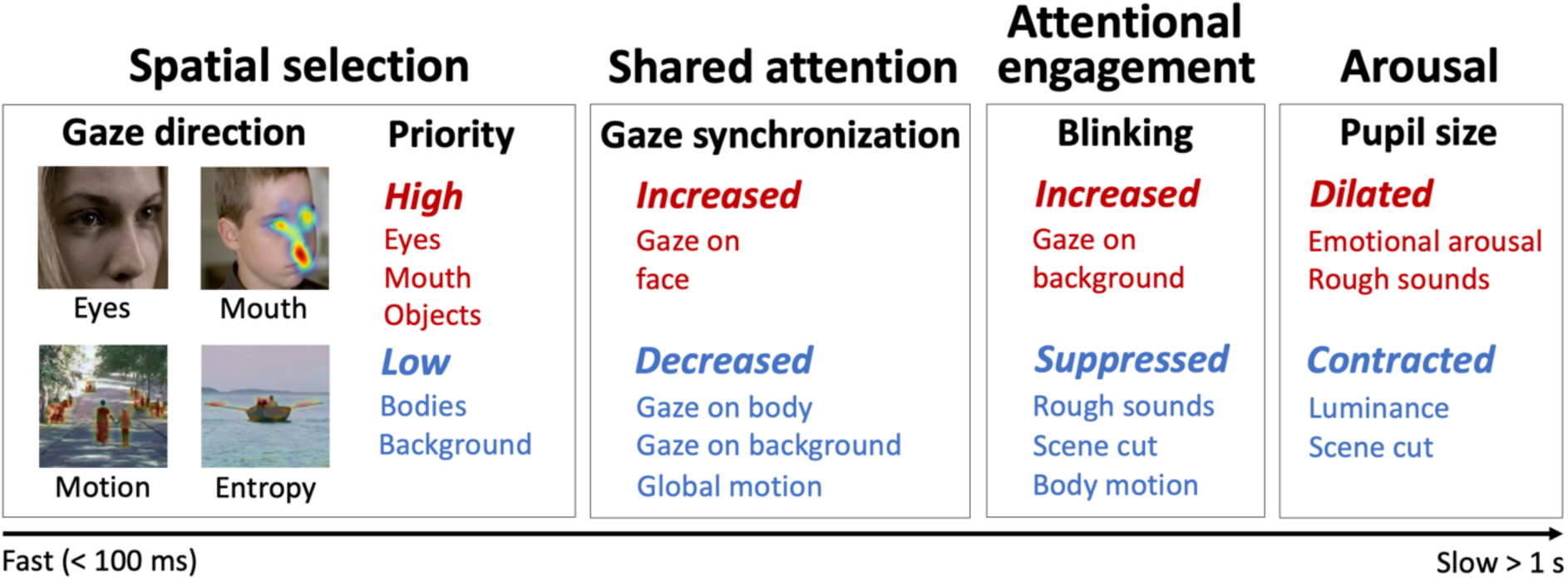
Summary of the findings. Investigation of the gaze direction, semantic category priority and gaze synchronization reveals that the presence of human faces, especially eyes, drive the fast visual sampling simultaneously with low-level visual features. Blinking is suppressed in intense scenes and during scene transitions indicating attentional engagement to the scenes, while pupil size is modulated by emotional arousal, scene transitions and luminance. The findings are based on analyses with 16 features derived from the movie stimuli (16 feature clusters were identified from the extracted 39 features). The sample images are from the Experiment 2 movie stimuli (Louhimies, 2008).

### Gaze direction depends on the presence of faces, motion, and visual information

The regression analysis for ISC and gaze prediction analysis both achieved moderate to high out-of-sample prediction accuracies (up to r = 0.43 for ISC and up to r = 0.47 for gaze heatmaps) indicating that external features consistently predict what we attend to in dynamic social scenes. Across the analyses we found that the eyes and mouths of the characters were the best predictors of dynamic gaze positions and the synchronization of gaze between participants. Across all datasets, people fixated on human eyes and mouth areas much more often than would be expected by chance (Figure 3). The developed permutation analysis ensured that the attentional priority for faces was not due to a centrality bias (Dorr et al., 2010) or just a result of movies presenting faces disproportionately more often than other contents. High priority for faces resulted in less fixations on bodies of the characters and on the background / surroundings, which were gazed much less than expected by chance. These results were confirmed in regression analyses that also accounted for low-level audiovisual features and perceived social information of the stimuli.

Eye movement synchronization (ISC) increased when faces were gazed at and decreased when the subjects gazed at bodies or background (Figure 4). This increased synchrony indicates that the faces, as biologically salient signal, could constitute a cue whose presence captures the viewers’ attention (Morrisey et al., 2019), thus resulting in increased gaze position synchronization. Indeed, the population gaze prediction analysis showed that eyes and mouth were the most important predictors of gaze positions in cinematic social scenes, surpassing low-level salient features (Figure 6). This is in line with previous findings suggesting that social information is prioritized over low-level saliency in social scenes (End & Gamer, 2017). Selective attention to human faces has been well established in controlled study designs (Bindemann et al., 2005; Morrisey et al., 2019; Theeuwes & Van der Stigchel, 2006). Other people are noticed quickly and independently of the task indicating that the identification of our conspecifics happens automatically (Rösler et al., 2017; Smith & Mital, 2013). Our results provide novel evidence on how faces are prioritized over other available information and provide an estimated “effect size” of the face priority when watching dynamic social scenes. The participants watched the faces 38 % (Exp. 1), 50 % (Exp. 2) and 38% (Exp. 3) of the total stimulus presentation time when only 14 %, 20 % and 10 % gaze times would have been expected if participants inspected the scenes with their eyes randomly.

In addition to social semantic features, local visual luminance / entropy predicted the moment-to-moment gaze positions (Figure 6). This visual feature cluster combined the information of “luminance” (a measure of local lighting), “visual entropy” (a measure of local randomness of pixel intensities) and “spatial energy” (a measure that detects local edges). Therefore, it indexes the amount of local visual information that the area conveys, as visual information requires light (luminance), local variation (entropy) and detectable forms (visual energy) to stand out from the background. High local contrast and low correlation between local pixel intensities (as measures of local information) have been previously shown to draw attention in natural still images (Krieger et al., 2000; Reinagel & Zador, 1999) supporting the interpretation that information-rich areas stand out from the rest and capture attention effectively.

While human faces were the strongest predictors of ISC and moment-to-moment gaze positions, *global* scene motion (Figure 4 **& Figure SI-4**) was associated with lower ISC indicating that the overall motion desynchronized the gaze positions between participants. The decrease in synchronization was most likely due to the need for increased visual sampling which resulted in diverging temporal gaze patterns between participants in these highly dynamic scenes. Nevertheless, the gaze prediction analysis indicated that areas with *local* motion draw attention (Figure 6). Increased attention to motion has been previously reported in humans (Abrams & Christ, 2003; Bruckert et al., 2023) and also in macaques (Mahapatra et al., 2008), and motion has been shown to capture attention task-independently suggesting automated attention towards motion (Smith & Mital, 2013). In sum, scenes with a lot of movement (real movement or camera movement) result in generally more scattered gaze positions between participants but local areas with high movement draw attention irrespective of the global amount of movement.

Scene cut effect analysis indicated that ISC increased briefly starting 200 ms after the scene cut and, depending on the stimulus, it returned to baseline between 800 ms (full movie stimulus) to 1400 ms (movie clips) after the scene cut (Figure 7). Higher ISC after scene transition suggests that introduction of a new scene leads to an orientation response that is tightly time-locked across subjects, while after this initial response the interindividual variation in the visual sampling increases. This accords with a prior study investigating the effect of scene cuts on gaze position synchronization (Mital et al., 2011). However, the regression analysis did not indicate a significant association between ISC and scene transition, although a consistent positive association was apparent in multiple different time windows (**Figure SI-4**). Since the regression results were carefully controlled for other predictors unlike the scene cut effect analysis, it is most likely that the observed increase in ISC after a scene change was more accurately modelled with other features, such as the presence of human faces.

Finally, fixation rate was higher when the participants gazed at semantic categories compared to unknown areas or areas outside the presentation window (Figure 4). Scene transitions were also associated with increased number of fixations but during intense and rough sounds the number of fixations decreased. Fewer saccades (decreased fixation rate) have been found to occur during high visual attention and with stationary stimuli (Mahanama et al., 2022). Hence, our results indicate that recognizing and tracking semantic objects and orienting to new scenes require more fixations (and saccades) compared to stationary moments with more static information. Intense and possibly emotionally arousing scenes are most likely visually engaging and saccades are inhibited to closely monitor the most important source of information.

### Pupil size depends on low-level information, emotional arousal, and scene changes

The multiple regression analysis for pupil size achieved high out-of-sample prediction performance (up to r = 0.5) indicating a major influence of external stimulus features on pupil size. In the regression models, the low-level feature luminance / entropy (including “luminance”, “visual entropy” and “spatial energy”) was negatively associated with pupil size, as the main function of the pupil is to regulate how much light enters the eye (Figure 4 **& Figure SI-4**). Perceived unpleasantness of the scene was positively associated with pupil size and this effect was independent from the overall scene luminosity. The unpleasantness predictor was combined from the evaluated features “unpleasantness”, ”aroused”, “aggressive”, and “pain”, based on the clustering analysis. This result indicates that unpleasant scenes were highly arousing in the movie stimuli. Additionally, we found that audio intensity and roughness was positively associated with pupil size. Intense sounds are arousing and alerting (Dean et al., 2011; Di Stefano & Spence, 2022; Ilie & Thompson, 2006; Trevor et al., 2020), which was also supported by the moderately positive association between perceived arousal and measured audio intensity (r = 0.39, **Figure SI-5**). The observed effect of emotional arousal on pupil size is in line with the previous controlled studies showing that both highly unpleasant and pleasant conditions dilate the pupil compared to neutral conditions (Bradley et al., 2008; R. R. Henderson et al., 2018; Partala & Surakka, 2003). Some studies also indicate that negative emotions may show a more robust effect on pupil size than positive emotions (Babiker et al., 2013; Kawai et al., 2013) – a result consistent with the present study.

The analysis of scene transition dynamics (Figure 7) showed that the pupil begins to contract rapidly after a scene transition with the peak contraction reached between 500 ms – 800 ms after the cut. Pupil dilates back to the baseline between 1150 ms – 1500 ms after the scene transition. This effect occurred with the same temporal scale as the ISC response to scene transition. Importantly, the regression analysis that controlled for other information also found the negative association between pupillary response and scene transition. Pupil constriction at the scene transitions may eventually reflect a more general response for quick change in the visual input, since the same phenomenon is found with controlled and simple stimulus changes under isoluminous conditions (Kimura et al., 2014). Nevertheless, the scene transitions should be taken into consideration in future studies using cinematic stimuli.

### Blinking as an indicator of attentional engagement

Participants lost only a few percent (median 2%) of the total stimulus viewing time due to blinking. Fewer participants blinked at the very beginning of a new scene (< 400 ms from the scene cut) compared to the later time points. The increase in ISC (and pupillary response) after scene transition begun simultaneously with the inhibition of blinking, providing further evidence that when a scene changes people really attend to the most important / interesting stimuli first and only afterwards they can blink without the risk of losing visual information (Figure 7). In line with this, we also observed a weak but consistent negative correlation (500ms scale: r = -0.14) between blinking and ISC when analyzing the full time series of the experiments (Figure 2). In the regression analysis, we did not observe a significant association between participant specific blink rates and scene cuts (Figure 4). People blink relatively infrequently when watching movies, around once per 5 seconds in our data, and this sparsity of blinks over the experimental time course most likely explains why significant associations between participant specific blink rate and scene change were not observed in the regression analysis. However, the blink rates were negatively associated with audio intensity / rough sounds and perceived body movement, which can be indicative of behaviourally intense moments in the scenes. This is also suggested by the relatively high correlations between arousal, aggression, body movement, auditory roughness, and audio intensity in the movie stimuli (**Figure SI-5**). People also blinked more often when they were gazing at the background, which could indicate that blinks occur during scenes with fewer important features, such as faces. In contrast with other modelled eye parameters, the regression models were poor at predicting the out-of-sample blink rates, suggesting that changes in blink rates may be more intrinsic than stimulus driven.

All in all, the results suggest that attentional engagement modulate the blinking behaviour. Previous studies focusing on blinks have reported that blinking is inhibited during attentional engagement (Ranti et al., 2020; Shin et al., 2015), blinking decreases as a function of attentional demand (Oksama & Hyönä, 2016) and blinks tend to occur at attentional breakpoints (Nakano et al., 2009; Wyly et al., 2024) supporting the current results derived from dynamic modelling of complex social scenes. Video stimulus with a coherent story line synchronizes blinking and yields lower blink rates compared to documentaries (Nakano et al., 2009; Shin et al., 2015), and blinking becomes increasingly synchronous across participants that are highly interested in the topic compared to those that are not (Nakano & Miyazaki, 2019). Functional brain imaging has demonstrated that the brain activity shifts from the dorsal attention network to the default-mode network after blink onset, which suggests that blinking associates with attentional disengagement (Nakano et al., 2013). Our results thus add evidence for the view that blinking reveals the participant’s attention or interest even dynamically when they watch complex social scenes, but that high-level social information has only negligible effects on blinking. Based on these confirming results, synchronized blinks could be used as indicators of attentional breakpoints of the stimulus and periods without (synchronized) blinks would indicate attentional engagement.

### Intense and rough sounds as an indirect measure of emotional arousal

The combined audio intensity and roughness was found to be a consistent modulator of pupil size, fixation rate, and blink rate in movie scenes. Most likely interpretation is that rough sounds indirectly capture intense periods in the movie stimuli, leading to people paying close attention to these events (suppressed blinking & lower fixation rate) and becoming emotionally aroused (pupil dilation). The audio intensity and roughness are physical properties that are easily extracted from audio stream and can be thus incorporated in any temporal analysis scale. Perceptual measures, such as evaluated emotional arousal, are subjective and the temporal analysis resolution depends on how the perceptual ratings are collected. Hence, audio intensity and roughness could serve as an indirect and objective measure of the behavioural intensity of movie scenes and emotion arousal of perceivers. Similar findings demonstrating that the auditory roughness relates to emotionally arousing (negative) events have been reported (Dean et al., 2011; Di Stefano & Spence, 2022; Ilie & Thompson, 2006; Trevor et al., 2020).

### Limitations

The stimulus contained short, socially rich Hollywood movie clips, which do not entirely reflect real world social situations. Movies have been intentionally directed and edited to capture attention and with editing and camera work the viewers’ visual attention is externally modulated in a way that diverges from how visual attention operates in real life. However, movies have the advantage of being controllable dynamic stimuli in the laboratory, since accurate eye tracking still requires a stationary measurement protocol. Viewing movies is a form of passive observation, while natural vision is used for guiding actions as well as for both gathering information from others and signaling back to them (Evan F. Risko et al., 2016). This dual function of gaze differentiates gaze patterns in real world situations from those in laboratory settings (Gobel et al., 2015; E. F. Risko & Kingstone, 2011). Future studies could collect first-person footage of every day social interaction with wearable cameras, while measuring the participants’ eye movements with accurate wearable eye trackers.

The initial feature space for the stimulus models was designed by the researchers. Hence, the results do not rule out the possibility that some other low-, mid- or high-level perceptual information can have significant influence on visual attention.

Additionally, the time-window approach taken in the current analyses treated consecutive time windows as independent samples. Since the consecutive samples in the time windows are never truly independent, even more detailed information could be extracted from the data, if temporal pattern information over multiple time windows were also analyzed. Future studies could investigate whether the social context could still yield specific temporal gaze patterns with dynamic stimuli. Finally, the results delineate the shared external influences of the visual system especially in social contexts. In doing so, they describe the average associations between external influences and human visual attention over individuals. In other words, they do not reveal any participant-specific factors or between-participant differences in how people pay visual attention in social contexts. Future studies on the attentional differences between individuals are needed for understanding how well the current models could predict the visual attention of certain individuals.

## Conclusion

Our results yield a comprehensive model on how visual attention, pupillary response, and blinking behavior are influenced by low-, mid-, and high-level perceptual features in dynamic perception using cinematic stimuli. We demonstrate that gaze, pupillary response and blinking behaviour are uniquely modulated by external stimulus features. Human faces, especially eyes, and low-level information (motion and visual information density) guide the gaze, not the socioemotional context. Pupillary response, in turn, was modulated by low-level information (mostly luminance) and emotional arousal, while fewer blinks occurred after scene transitions and during behaviourally intense scenes (loud noises, body movement). This work advances our understanding of visual processing in social contexts by modelling different eye-tracking parameters dynamically and simultaneously. Future research should explore how visual attention is modulated in even more complex and ecologically valid environments to bridge the gap between controlled laboratory studies and fully natural perception.

## Materials and Methods

### Experimental design

To investigate how the visual perceptual system works in dynamic social contexts, we set up three independent eye-tracking experiments, where participants watched different movie stimuli rich in social interaction. Different participants were recruited for each experiment. In Experiment 1, we used our previously validated socioemotional “localizer” paradigm that allows us to present variable social content (Karjalainen et al., 2017, 2018; Lahnakoski et al., 2012; Nummenmaa et al., 2021; Santavirta et al., 2023). The experimental design and stimulus selection has been described in detail in the original study with this setup (Lahnakoski et al., 2012). Briefly, the participants viewed a set of 68 movie clips (median duration: 11.2s, range: 5.3 – 27.8 s, total duration: 14 min 26 s) that have been curated to contain large variability of social and emotional content. The videos were extracted from mainstream Hollywood movies with audio tracks in English. See **Table SI-1** for short descriptions of the movie clips. In Experiment 2, the participants watched a Finnish historical drama film “Käsky” ((Louhimies, 2008), 70 min 14 s), and in Experiment 3, the participants watched a full-length horror movie “The Conjuring 2” ((Wan, 2016), 109 min 3 s). The total stimulus duration of the stimuli was 193 minutes 43 seconds in three independent experiments. The variability of the movie stimuli ensures that the replicable findings would generalize across different social perceptual contexts, at least within cinematic stimuli.

### Participants, eye-tracking data, and quality control

A total of 166 participants were recruited (Experiment 1: 110 participants, Experiment 2: 28 participants, Experiment 3: 28 participants) and the data were collected in Turku, Finland between 2019 and 2021. Normal or corrected to normal vision was required. Participants gave an informed consent prior to taking part in the experiment. Seven participants were excluded due to incomplete data (three for incomplete data and four for corrupted data files resulting in partial data loss). We also excluded participants from the analyses based on data-driven quality control of the eye-tracking data. We first calculated the total time of fixations and blinks as well as the gaze position time within the presented video area (time in presentation area / total experiment duration) for each participant. Then we fitted beta distributions to these data and identified the 2% probability cutoff points from the long tail of the fitted distributions (**Figure SI-6**). The participants whose total fixation time, total blink time or total gaze time fell into the 2% probability in the long tail of the distribution were considered outliers and their data were excluded. Four participants were excluded because they had both atypically low total fixation times and high blink durations (fixations less than 76% and blinks over 12% of the total time), two participants based on low fixation time alone (fixations less than 76% of the total time) and two participants based on low total time within video area (gaze position within video area less than 93% of the time). The final sample thus included 151 out of 166 participants.

### Eye movement recordings and data preprocessing

In Experiment 1, short movie clips whose contents were unrelated to each other were presented to the participants in fixed, initially randomized order without breaks to enable synchronization analyses. The eye tracker was calibrated and validated using a 5-point calibration, and validation was repeated with one point before the experiment. Validation was successful if gaze position error was below 1°. Validation was repeated three times during the experiment (after every 17^th^ movie clip). The stimuli were presented with a 27’’ Retina 5K monitor at 90 cm distance from the eyes. The eye-tracking data was collected with SR EyeLink 1000 Plus (SR Research, Ontario, Canada) eye tracker with the following setup: v5.15 Jan 24 2018, Eyes: Right, File filtering level: Extra, Pupil tracking algorithm: Centroid. In Experiments 2 and 3, the movie was presented in short segments in fixed chronological order (Experiment 2: 26 trials, median trial duration: 159 s, range: 68 s – 256 s. Experiment 3: 30 trials, median trial duration: 215 s, range: 146 – 300 s) to allow repeated drift correction and camera calibration throughout the experiment and to maximize participant comfort. Before each trial (i.e. movie segment) a 5-point calibration and validation and a subsequent 1-point validation (error < 1°) was conducted. The stimuli were presented with a 24’’ BenQ XL2411Z monitor at 70 cm distance from the eyes. The eye-tracking data were collected with EyeLink 1000 (SR Research, Ontario, Canada) eye tracker with the following setup: v4.594 Jul 6 2012, Eyes: Right, File filtering level: Extra, Pupil tracking algorithm: Ellipse. Fixation and saccade reports were generated with EyeLink DataViewer 4.1.1. software (https://www.sr-research.com/data-viewer/). Fixations shorter than 80 ms were considered unreliable and previous reliable fixation was extrapolated to continue until the next reliable fixation to create a continuous time series of fixation information.

### Eye-tracking parameters

We extracted pupil size, gaze position, saccades, and blink timings from the EyeLink DataViewer reports. Intersubject correlation of the gaze position (ISC) was dynamically calculated for measuring the moment-to-moment gaze position synchronization across participants using eISC-toolbox for Matlab (Nummenmaa et al., 2014). Briefly, the ISC is based on computing participant-wise fixation heatmaps in pre-defined time windows, and momentary ISC is defined as the mean spatial correlation across participants in each window. For gaze prediction analysis (see section “Gaze prediction analysis”), population average gaze heatmaps were also generated using the eISC-toolbox. Blink synchronization was estimated by calculating how many participants blinked within a given time window. Fixation and blink rates were calculated separately for each participant in specific time windows.

A single analysis time window such as 500 ms or 1000 ms is not justifiable in uncontrolled movie perception because different eye movement parameters (e.g. pupillary responses or gaze position changes) have different intrinsic time scales, and because the predictor variables also had different time scales (from 40ms resolution for physical and semantic features to 4000 ms resolution for the perceived social features, see “Perceptual models derived from movie stimulus”). Therefore, different analysis time scales may capture associations between the visual system and the stimulus features. To estimate the effect of the chosen temporal scale, the eye-tracking parameters were sampled and analyzed across different temporal scales (200 ms, 500 ms, 1000 ms, 2000 ms, 4000 ms).

### Perceptual models derived from the movie stimuli

Stimulus features varying from low-level audiovisual properties (e.g. luminance and audio intensity) to semantic information (e.g. faces and objects) and to the perceived socioemotional information (e.g. (un)pleasant interaction and arousal) were estimated from the video stimuli to model the simultaneous effects of perceived features across distinct processing levels in human cognition. We 1) extracted multiple low-level features and their time derivatives from the audiovisual domain (24 features), 2) identified semantic features from the video frames using computer vision (7 features) and 3) collected dynamic perceptual ratings of high-level socioemotional features from human annotators (8 features). The predictors were then sampled to different temporal scales (200 ms, 500 ms, 1000 ms, 2000 ms, 4000 ms) matching with the eye-tracking data (see above).

### Low-level audiovisual features

Low-level visual features were extracted for each frame (static features) or consecutive frame pair (features that measure change) from the video stimulus. Visual features included luminance, visual entropy, optic flow, spatial energy with two distinct Fourier filters for edge detection, and differential energy for measuring total change between consecutive frames. Auditory features included audio intensity (RMS), properties of the frequency spectrum (geometric mean, standard deviation, entropy, and high-frequency energy), waveform sign change rate or “noisiness” and sensory dissonance or “roughness”. The auditory features were extracted using MIRToolbox (Lartillot & Toiviainen, 2007). Since the video frame rate was 25 / second, the auditory features were extracted in matching 40 ms time windows. For each audiovisual feature (except optical flow and differential energy that are already measured between frames) the difference between consecutive frames was also calculated. The visual information was extracted locally within each frame and then averaged over the pixels in each frame. Detailed properties of the feature extraction are provided in Supplementary materials (see section “Low-level feature extraction”).

### Semantic features

We segmented the stimulus films frame-by-frame to extract semantic information from the videos using pre-trained open-source computer vision models. Each pixel was assigned into a class (eye area, mouth area, other face area, body parts, animals, objects or background) or left unknown, if the model’s prediction confidence was under

50 %. To first segment the whole image, we used a panoptic feature pyramid network (FPN) segmentation model (https://github.com/facebookresearch/detectron2/blob/main/configs/COCO-PanopticSegmentation/panoptic_fpn_R_101_3x.yaml) from the Deceptron2 Python library (Wu et al., 2019). The FPN included a Mask R-CNN architecture with ImageNet pretrained ResNet101 backbone and it was trained on the COCO dataset to segment the images to 134 initial categories (Kirillov et al., 2019; Lin et al., 2014). After segmentation, the initial categories were assigned into broad semantic classes (bodies, animals, objects, background and unknown) for the analyses. Next, we used the RetinaFace face detection model (Deng et al., 2020) following the implementation of a previous eye-tracking study of autism to segment rectangular face, eye and mouth areas from the videos (Keles et al., 2022). Eye and mouth areas were excluded from the whole face segmentations to parcel faces into three segments (eyes, mouth and face excluding eyes and mouth).

The accuracy of the automatic segmentations was validated using human reference in the stimulus for Experiment 1. Judging the accuracy of all segmentations from each stimulus frame would have been overly laborious. Hence, the human annotator evaluated the accuracy of the segmentations in the most interesting areas (i.e. areas receiving most fixations from observers) of a subset of the video frames as follows. One frame per second was extracted from the stimulus and the segmentations for this frame were overlaid onto the frame. One second interval was considered sufficiently short for accurate estimations of the segmentation data quality and the work was not overly laborious for manual checking with that interval. Next, the thresholded (95^th^ percentile) population average gaze position heatmap was overlaid to the frame to identify the areas that were the most interesting to the participants around the extracted frame (100 ms time window around the frame). A human annotator checked these images one-by-one and judged whether the segmented classes within the interest areas (under the gaze position heatmap) were correct or not, and then marked the right class based on her opinion. The model’s average positive predictive values were: eye area 99 %, mouth area 87 %, other face area 99 %, bodies 69 %, animals 65 %, objects 79 % and background 83 %. The sensitivities were: eye area 92 %, mouth area 95 %, other face area 77 %, body parts 89 %, animals 59 %, objects 44 % and background 82 %. The confusion matrix of the quality control results, and sample frames of the segmentations are shown in Figure 9. Since animals were infrequently present and the positive predictive value of this category was the lowest, we decided not to include this category in the further analyses.

**Figure 9.**
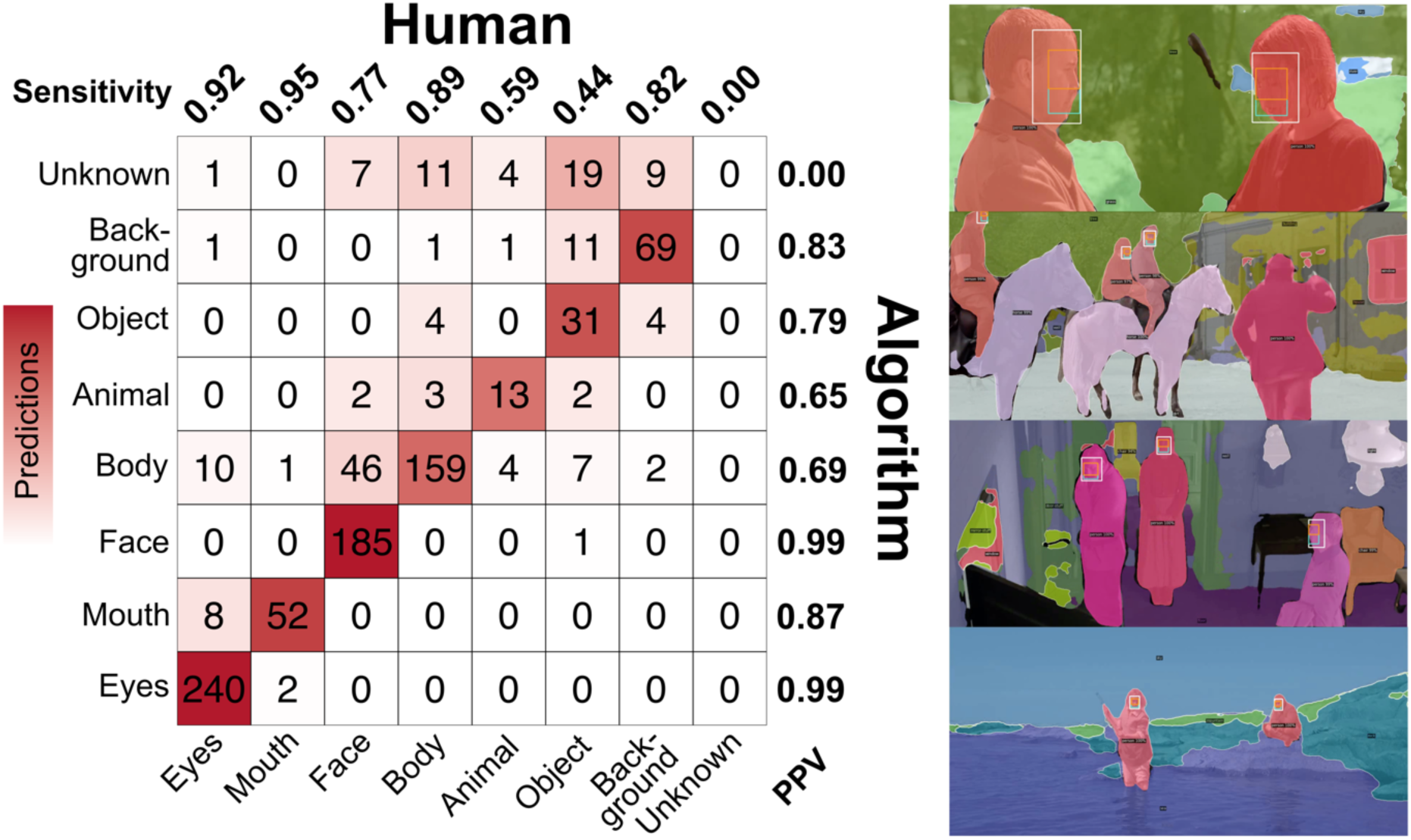
Semantic segmentation quality control. The confusion matrix shows the agreement of a human annotator and the segmentation algorithm in the Experiment 1 (movie clips) stimulus. Examples of the segmentation results are visualized alongside the confusion matrix. The sample images are from the Experiment 2 movie stimuli (Louhimies, 2008).

### Scene cuts

Unlike real life situations, movies contain scene cuts that result in major semantic and perceptual changes. The cuts also impact eye movements considerably (Bruckert et al., 2023). We used a scene filter from ffmpeg to detect scene changes in the stimulus (https://ffmpeg.org/ffmpeg-filters.html#select_002c-aselect, filter command: “select=’gt(scene,threshold)’”). The filter first assigns a score to each frame, where higher scores indicate higher probability of the frame starting a new scene. Cuts are then identified using a detection threshold for the score. To optimize the detection threshold and validate the performance of the automatic scene cut detection method, three human annotators identified the scene cuts in the Experiment 1 stimulus. Using the manual annotations as the ground truth, the detection threshold of the automatic scene filter was optimized to balance the positive predictive value and sensitivity of the scene cut detection. Two hundred ms temporal distance between the algorithm and human annotators was chosen as the maximum distance for confirming a match. Optimized scene cut detection achieved 96% positive predictive value (PPV) and 95% sensitivity in the Experiment 1 stimulus. The optimized scene filter was used to detect scene cuts in all three experiment stimuli.

### Perceived social features

We selected a set of perceived socioemotional features that were identified as important perceptual features in our previous studies of social perception (Santavirta et al., 2023, 2024). These features are also annotated with high between-participant consistency (Santavirta et al., 2024) which is crucial since different participants annotated the stimuli and participated in the eye-tracking experiments. These features were perceived pleasant feelings, unpleasant feelings, arousal, pain, talking, body movement, feeding and playfulness. The ratings were collected in four-second temporal resolution with an abstract slider scale from “absent” to “very much”. Five annotators were recruited to the laboratory for annotating these features for Experiment 1 and 2 using Onni platform (https://onni.utu.fi/) (Heikkilä et al., 2021). The annotations for the Experiment 3 were collected online using Gorilla platform (https://gorilla.sc/) and the participants were recruited through Prolific (https://www.prolific.com/). To decrease the workload of online participants one participant annotated only a subset of the video clips. The online experiment begun with a bot check (in addition to Prolific’s internal bot checks for their participant pool) to increase the data quality. Quality control of the online participant data was conducted as described previously (Santavirta et al., 2024) and data were collected until five reliable annotations were retrieved for each clip and feature. Finally, we took the average over the individual annotators to get the high-level predictors for the eye-tracking analyses.

### Final predictor set

As described in above sections, we extracted a total of 39 perceptual features from the video stimulus in order to get a detailed description of the stimulus contents in multiple processing levels from pure audiovisual processing to semantic categorization and perceived social information. Correlation analysis between the extracted features indicated that some of the features were strongly associated with each other. Reliable and interpretable mapping of the eye-tracking data with the available features requires that the predictors provide relatively independent information. We thus decreased the correlations between predictors by hierarchically clustering the correlation matrix of the extracted features using the default UPGMA clustering algorithm (https://www.rdocumentation.org/packages/stats/versions/3.6.2/topics/hclust). The clustering identified 16 clusters (seven clusters for low-level features, five clusters for perceptual semantic categories and four clusters for perceived social information) from the extracted 39 features which were used as predictors in the analyses instead of the original features. Cluster predictors were formed by calculating the average over the standardized values of the features within each cluster. See section “Dimension reduction” and **Figure SI-5** in the supplementary materials for a detailed description of the dimension reduction.

### Statistical analyses

The aim of this project was to establish how external low-, mid-, and high-level social perceptual features together guide the visual system. The following research questions were studied with unique analytical techniques developed for each of them:

1. Do people prioritize certain semantic categories (e.g. faces) over other semantic categories in social scenes?
2. How are the eye-tracking parameters dynamically modulated by low-level audiovisual information, semantic category information, and perceived social information in social scenes?
3. How accurately the population level gaze directions can be dynamically predicted in social scenes with an interpretable model that incorporates low-level information, semantic category information, and perceived social information?
4. How scene cuts influence the eye-tracking parameter dynamics immediately after a scene cut?

To address each of these questions, we developed a set of analytical methods and tested whether the results generalize across the three independent datasets. First, we conducted a gaze time analysis to assess if people have attentional priorities for some semantic categories (**Question 1**). Second, we used a cross-validated multiple regression analysis to assess which perceptual predictors have an independent association with pupil size, ISC, fixation rates and blink rates (**Question 2**). Third, we built cross-validated random forest models for moment-to-moment pixelwise predictions of the population level gaze heatmaps to understand how well the actual gaze positions can be modeled with the available predictors and to deepen the understanding of the attentional priorities when observing dynamic social scenes (**Question 3**). Finally, we investigated the average dynamics of the pupil size, ISC and blink rates after each scene cut (**Question 4**).

### Gaze time analysis

To investigate whether people have an attentional preference towards certain semantic classes we conducted a gaze time analysis. The pixel-to-pixel semantic class information from the computer vision allowed identifying which class (eyes, mouth, face excluding eyes and mouth, body parts, object, or background) a participant was watching at each time point. From these data, we calculated the total gaze time for each class and averaged the gaze times over the participants within each experiment to get population level gaze times for each class. The analysis was conducted separately for each experiment.

High gaze time for a class does not itself reveal an attentional preference for the class but could merely indicate that the class is often present occupying large parts of the screen. Additionally, centrally presented contents are likely to receive more fixations as observers prefer looking at the center of the stimulus (Dorr et al., 2010). To control for the location, frequency, and total presented area for the semantic classes in our stimulus we used permutation testing to differentiate whether the true gaze times are significantly different from those expected by chance. The permutation testing was conducted by bootstrapping the true fixation gaze data circularly to break the temporal synchrony between the stimulus and the gaze coordinates. Total gaze times for semantic classes were then calculated for the bootstrapped random data. The bootstrapping was repeated 500 times to get the estimation of the gaze time null distributions. Finally, a p-value for the null hypothesis that the true gaze time does not differ from chance was calculated by ranking the real gaze times to the corresponding null distributions. Since true fixation coordinates were used in the permutation testing instead of simulated random coordinates, the only difference between random data and true data is the temporal asynchrony between the gaze coordinates and the stimuli (distributions of gaze coordinates and fixation durations do not differ).

### Regression analysis

To investigate the associations between the extracted predictors and eye-tracking parameters while controlling for all other predictors we conducted a multi-step regression analysis. Even after dimension reduction the 16 features in the predictor space were not fully independent. Repeated simple regressions or one multiple regression with all the predictors in the same model could result in mixed findings due to possible collinearity between predictors. Instead, we chose a multi-step regression approach to achieve more support that the identified associations are truly independent of the effects of other included predictors. For pupil size modelling, predictors were shifted 1000 ms forward to correct for the lag in pupillary response (**Figure SI-7**) while ISC, fixation rate and blink rate were modelled without temporal shift.

First, we ran simple regression for all predictors in a leave-one-experiment-out cross-validation process where models were fit to the data of two experiments and one experiment’s data was left as the testing set. The predictors where the sign of the association (regression coefficient) was consistent across all cross-validations indicating replicable sign of association between experiments were selected for the next analysis step. If the simple regression associations were inconsistent between the cross-validation rounds, we concluded that the results do not support association between eye-tracking data and the feature. Second, we ranked the consistent predictors by their out-of-sample prediction accuracy (correlation of the model’s predictions with the leave-out experiment data) to prioritize the features that have higher predictive power in the simple regression setting over the features with lower predictive power. Finally, we built an additive regression model where we added candidate predictors one-by-one to a multiple regression design starting with the features whose predictive power ranked the highest in the previous regression model. After every addition of a new predictor to the model, we tested whether the out-of-sample prediction accuracy (correlation with the leave-out experiment data) improved more than would be expected by chance. If the prediction accuracy after adding a new predictor was lower than expected by chance, we concluded that the predictor does not have an independent association and the predictor was dropped from the design. Hence, the final model tests all consistent predictors and returns the final model with only predictors that improve the prediction accuracy significantly.

The order in which the predictors are added to the multiple regression influences the results when predictors are not fully independent but testing all possible combinations and orders is computationally prohibitive with up to 16 predictors. Hence, the initial prediction accuracy in the simple regression was chosen to determine the order of addition. This approach penalizes the predictors that initially showed a low prediction accuracy but does not rule out the possibility of an independent effect if the predictor’s effect is not better modelled by the previously added predictors. This approach assumes that if multiple correlated predictors had association with the variable of interest in the simple regression setup, the one with the highest prediction accuracy would be the one most likely associated with the variable of interest. To ensure that the simple regression and the stepwise multiple regression results did not yield mixed findings, we confirmed that the sign of association of the final set of significant predictors stayed the same in the initial simple regression and in the multiple regressions.

To test whether adding a predictor to the multiple regression design yields better predictions than expected by chance in the left-out-experiment data we conducted a permutation test after each predictor addition. Only the newly added column of the design matrix was bootstrapped circularly to preserve the covariance structure of the previously validated model. Bootstrapping was repeated 500 times to generate the null models. The null distribution of the out-of-sample prediction accuracy was generated from the prediction accuracies of these null models. The p-value for the hypothesis that the newly added feature did not improve the model’s prediction accuracy more than would be expected by chance was then calculated by ranking the true prediction accuracy to the null distribution. The predictor was considered to improve the prediction accuracy significantly when p < 0.05.

### Gaze prediction analysis

We built a data-driven model to predict the population average gaze position heatmaps to assess how well the current predictors can explain the momentary gaze directions of the subjects. Pixelwise predictor values were used to predict the gaze probability of each of the pixels. To lower computational demands, the gaze heatmaps and predictors were downsampled to 64 x 64 pixel resolution prior to training the gaze prediction models. We did not model the gaze heatmaps for each frame since 40 ms time interval is unnecessarily short. Instead, we calculated the gaze heatmaps in 200 ms time windows for this analysis to preserve high temporal resolution while allowing the temporal accumulation of information between a few adjacent frames. This temporal resolution is likely close to the temporal resolution of the human visual system since the typical fixation duration was approximately 300 ms in our data (Figure 1).

We used a random forest regression model that preserves some interpretability, allows for more complex association to be considered than ordinary linear regression and is computationally efficient, unlike deep convolutional neural network which would be computationally prohibitive and difficult to interpret in this context. The random forest models were trained using the fitrensemble function in Matlab (https://se.mathworks.com/help/stats/fitrensemble.html). The fitrensemble function randomly selects N out of N observations of data with replacement and ⅓ of predictors for training each individual decision tree. The number of individual decision trees and the number of branches in each tree were optimized (see supplementary materials, section “Random forest regression optimization”). Based on the optimization results 50 decision trees and 63 branches (6 branches from the tree trunk to the leaf) were selected. Separate models were trained for each of the three datasets to allow out-of-sample performance testing and comparison of the trained models between datasets.

The performance of the gaze prediction models was evaluated by testing the trained models on the two datasets that were not included in the model training. Two performance metrics were calculated. First, we calculated the correlation between the true heatmap values and the predicted heatmap values over all time windows in the tested dataset. Correlation measures the overall similarity of the heatmap distributions but is not sensitive in measuring how well the model predicts the most salient area (area with the peak heatmap value) in the screen. To assess how well the model can predict the most salient areas, we also calculated the distance between the true peak value and the predicted peak value for each 200 ms time window. This peak prediction distance was then averaged over all time windows and is reported as percentage of the image width. If the peak prediction distance is low, then the model is able to predict well the most salient part of the screen, regardless of the overall similarity of the true and predicted heatmap distributions.

To interpret the trained random forest models we extracted the relative feature importance (https://se.mathworks.com/help/stats/classreg.learning.classif.compactclassificatione <u>nsemble.predictorimportance.html</u>) for each predictor which states how influential the predictor was for the model’s prediction in range between 0 and 1. The relative importance does not reveal whether increase in the predictor value would increase the predicted gaze probability or vice versa. Hence, we simulated how changing the value of one predictor influences the model’s predictions when all other predictor values are held constant. First, we randomly selected a pixel from the training dataset and extracted the real predictor values for that pixel. For categorical predictors we simulated how changing the value between 0 and 1 will influence the gaze probability prediction in that pixel. For standardized continuous predictors we simulated new predictions with 20 uniformly distributed random predictor values (Z-scores) between -3 and 3. This simulation was repeated for 200 000 randomly selected pixels and the simulation was done separately for each trained model (one for each dataset) before pooling the simulation results together.

### Scene cut influence analysis

We extracted the eye-tracking data for 3600 ms time window around each identified scene cut (from cut – 600 ms to cut + 3000 ms) for each participant. Pupil size, ISC and blink rate were then extracted for these time windows. Pupil size was extracted continuously (1kHz) and ISC and blink rates were calculated in 200 ms time windows (5Hz). Since pupil size is an arbitrary participant specific measure, pupil size time series were normalized with the mean pupil size in a 600 ms interval before scene cut. To estimate the replicability of the scene cut effect, the data were averaged over the participants separately for each experiment.

To estimate whether the population level change in eye-tracking measures after scene cut was significantly higher than would be expected by change, a permutation test was applied. We sampled 100 time series (3600 ms) at random time points from the eye-tracking data for each participant and averaged these over all participants to get a random sample of the population level average time series for the 3600 ms time period. This sampling was repeated 500 times to get a null distribution of the population level random time series thus the null distribution was generated from 50 000 random samples (100 x 500). True population level time series calculated from participant-level time series around scene cuts were ranked within this null distribution to get the exact p-value for each time point around the scene cut. Significant cut effect is reported for time points with p < 0.05 based on this permutation test.

## Data and code availability

The analysis scripts and the extracted stimulus features are freely available in the project’s GitHub repository (https://github.com/santavis/social-vision-in-cinema). The subjects’ permissions for public distribution of the subject-level eye-tracking data were not collected and hence the eye-tracking data cannot be distributed. The stimulus movies can be made available for researchers upon request, but copyrights preclude public redistribution of the stimulus set. **Table SI-1** provides short descriptions of the movie clips in Experiment 1.

## Declaration of competing interest

The authors declare no competing financial or non-financial interests.

## CrediT authorship contribution statement

**Severi Santavirta**: Conceptualization, Methodology, Software, Validation, Formal analysis, Investigation, Resources, Data curation, Writing – original draft, Writing – review & editing, Visualization, Project administration. **Birgitta Paranko**: Conceptualization, Investigation, Resources, Data curation, Writing – review & editing. **Kerttu Seppälä**: Conceptualization, Investigation, Resources, Data curation, Writing – review & editing. **Jukka Hyönä**: Conceptualization, Methodology, Resources, Writing – review & editing. **Lauri Nummenmaa**: Conceptualization, Methodology, Resources, Writing – original draft, Writing – review & editing, Visualization, Supervision, Project administration.

## Acknowledgements

The study was supported by Turku University Foundation and Alfred Kordelin Foundation grants to SS and Finnish Governmental Research Funding for Turku University Hospital and for the Western Finland collaborative area to SS. We thank Elina Lewandowski for her help in the eye-tracking data collection and Maya Rassouli for the image segmentation quality control.

## Notes

https://github.com/santavis/social-vision-in-cinema

### Competing Interest Statement

The authors have declared no competing interest.

