## Supplementary materials for "Modelling human social vision with cinematic stimuli"

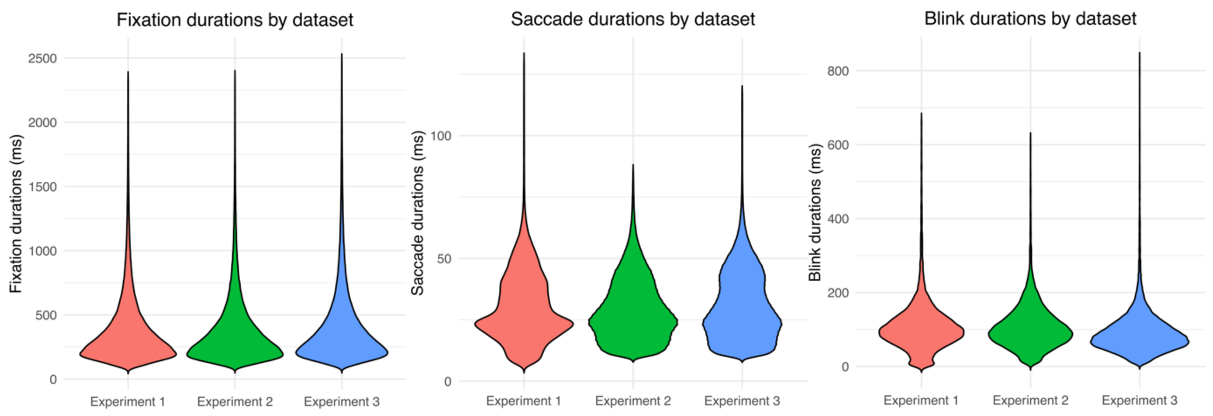


**Figure SI-1.** Fixation, saccade, and blink durations separately for each dataset.

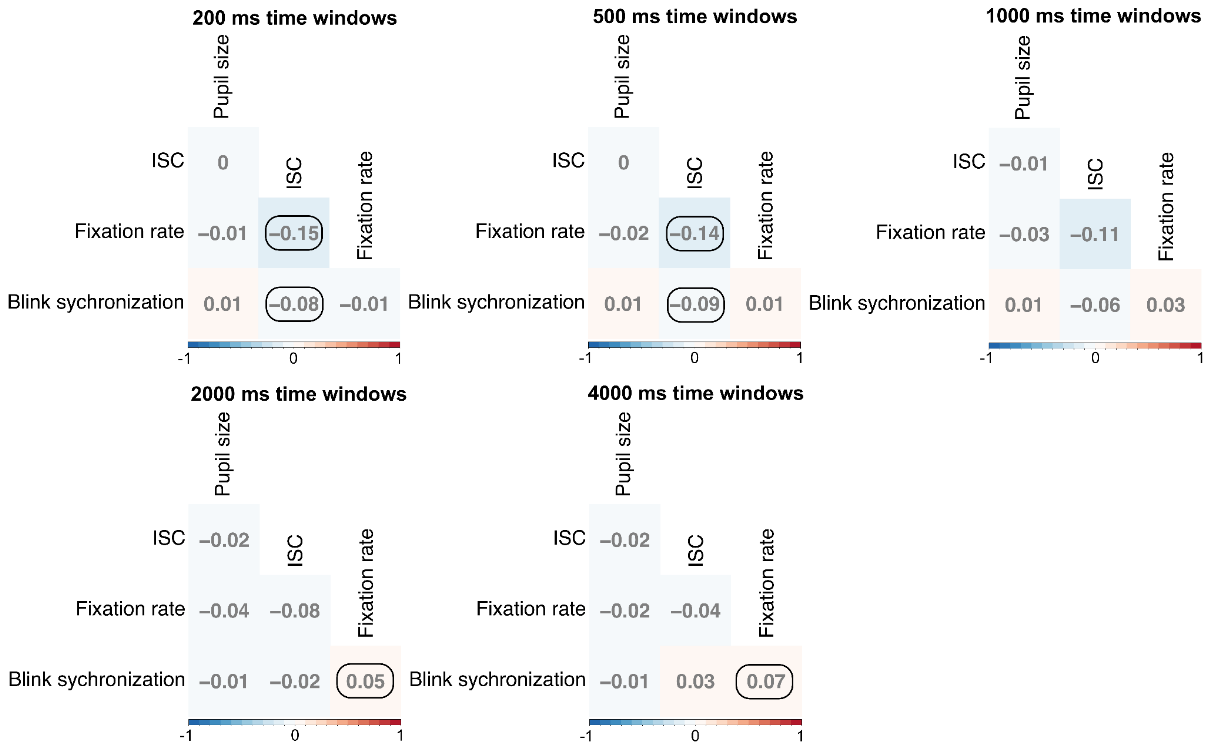


**Figure SI-2.** Correlations between the eye-tracking variables. The correlations were calculated for each experiment separately and then averaged over datasets for the visualizations. Correlations that had consistent signs over all experiments are circled. The correlations were calculated in multiple different time scales (200 ms – 4000 ms). Blink synchronization here measures how many participants blinked in a given time window.

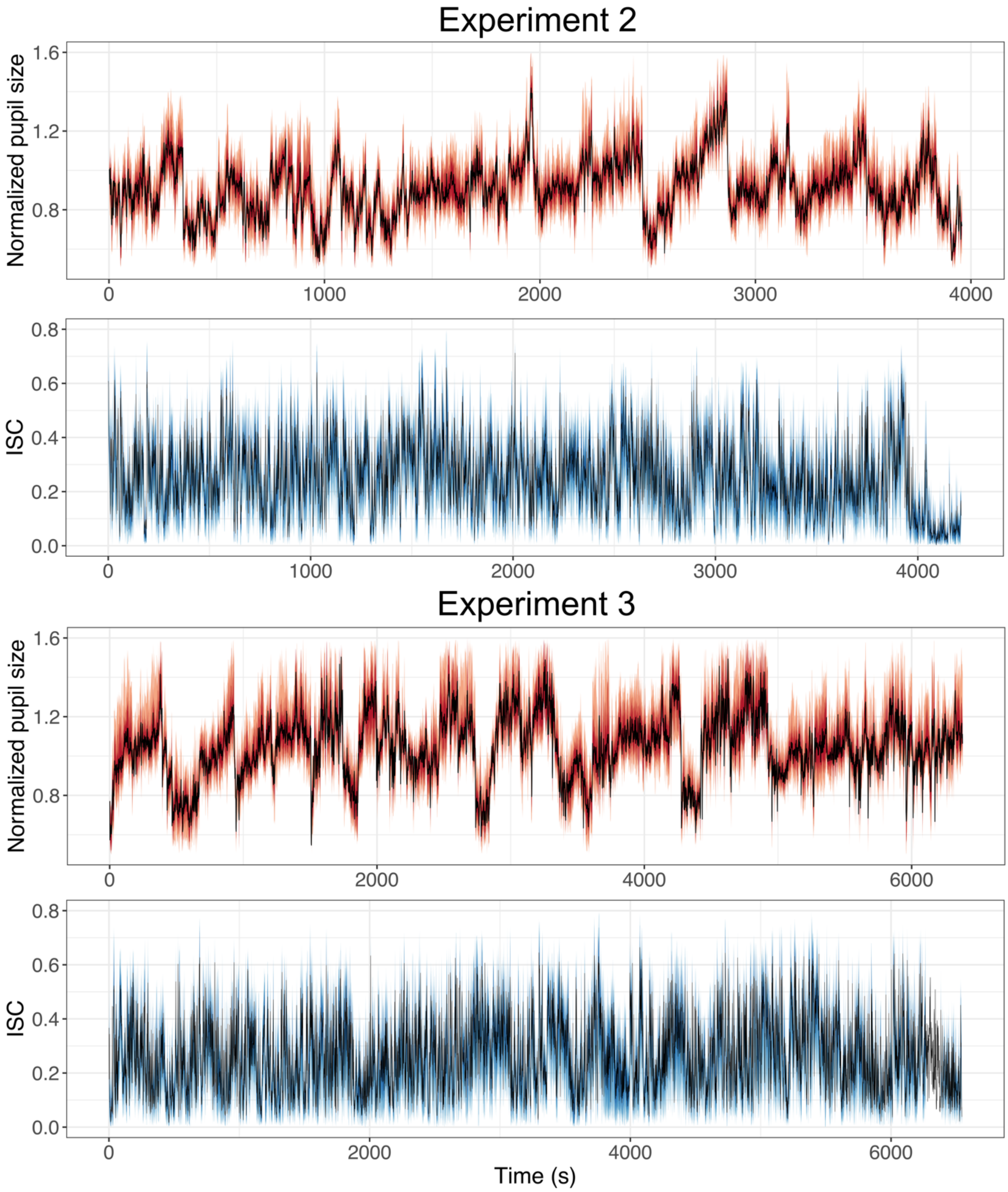


**Figure SI-3**. Time series of median pupil size and ISC across subjects in the Experiment 2 & 3 (coloured areas mark the 40%, 60% and 80% quantile intervals).

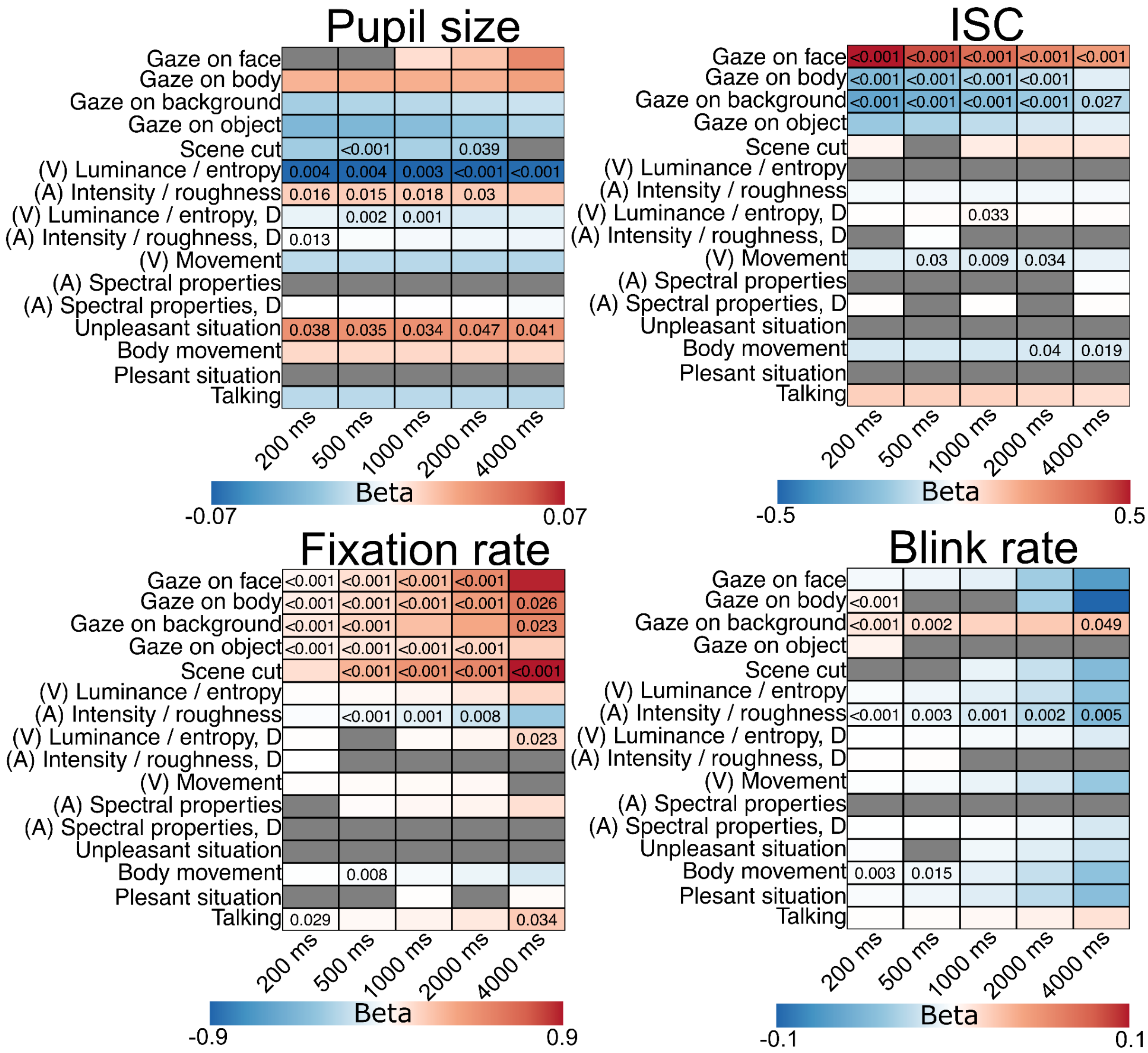


**Figure SI-4.** Regression results for all analyzed time scales. Pupil size, ISC, fixation rate and blink rate were analyzed in five different time scales (200ms – 4000ms) with the same set of (16) features. Positive associations are coloured in red and negative associations in blue. Grey colour indicates that the direction of association was inconsistent between the three datasets. The overlaid numbers visualize the permuted p-values of the statistically significant ($p < 0.05$) associations.

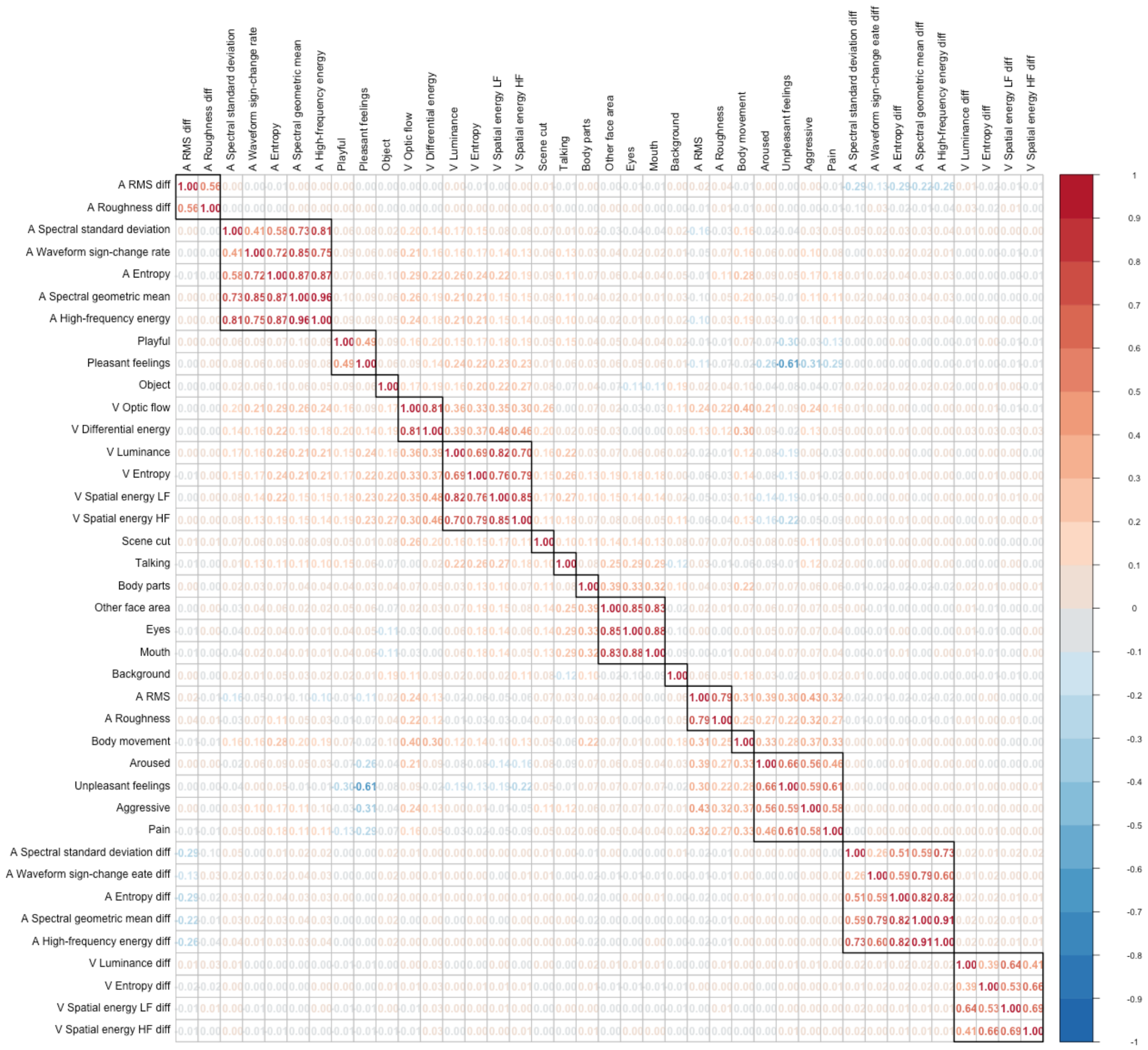


**Figure SI-5.** Clustered average correlation matrix of the extracted perceptual features over the three experiments and over the different analysis time windows.

### Low-level feature extraction

For better interpretability of the results, we extracted an extensive set of audiovisual physical features from the stimulus films instead of letting deep AI models define low-level features that may be hard to interpret. In the visual domain, we estimated luminance, entropy, and spatial energy for each frame. **Luminance** was estimated as the 3rd dimension (V, values) of the RGB to HSV transformation. **Visual entropy** was estimated from the grayscale image as $-\sum(p*\log_2(p))$ where p contains the normalized histogram counts (https://se.mathworks.com/help/images/ref/entropy.html). Spatial energy was calculated to detect edges in the image. For spatial energy calculation the images were Fourier transformed and low-pass filtered in the frequency domain to cut out high frequency noise. The corner frequencies (high frequencies in both X and Y axis) were filtered out as probable noise. By visual inspection of the Fourier filtered images two different Fourier filters were used to detect different frequency edges in the images. For feature **spatial energy HF**, corner frequencies within the radius of 1% of the Y resolution of the image were filtered out. The filter radius for the **spatial energy LF** was 10% of the Y resolution and therefore more corner frequencies were filtered out compared to the "Spatial energy HF". To measure movement and change between consecutive frames of the video **spatial energy** and **optic flow** were calculated. Spatial energy was calculated as the RMS of the pixel-to-pixel change between consecutive frames. Optic flow was calculated with Matlab's estimateFlow function (https://se.mathworks.com/help/vision/ref/opticalflowhs.estimateflow.html) using Lucas-Kanade method. For the features calculated for each frame (luminance, entropy, and spatial energy) the difference between consecutive frames was calculated to get detailed temporal understanding of the stimulus. The pixel specific values were used in the gaze prediction analysis, but averages over the framewise pixels were used in the regression analysis.

MIRToolbox (Lartillot & Toiviainen, 2007) was used to extract detailed auditory information of the stimulus. The auditory features were calculated in 40 ms time windows corresponding to the framerate of the videos (25 / s). **Audio intensity** was estimated as the RMS of the audio stream (mirrms function). To get a detailed understanding of the frequency spectrum, the **geometric mean**, **standard deviation**, **entropy,** and **high-frequency energy** (the cutoff where 85% of the energy is below it) were extracted using functions mircentroid, mirspread, mirentropy and mirrolloff, respectively. **Waveform sign change rate** as the estimate for "noisiness" was calculated with function mirzerocross. Sensory dissonance or "**roughness**" of the sound was calculated using function mirroughness. To estimate the change in the auditory properties, the change between consecutive time windows was calculated.

Dimension reduction

The extracted feature set contained 39 features (24 low-level audiovisual features, eight semantic features and seven perceived social features). Since some of the extracted 39 perceptual stimulus features were highly correlated (up to $r > 0.9$) we hierarchically clustered the correlation matrix of the predictors as a dimension reduction method. To investigate the predictor correlation structure on the population level, perceived semantic features (what the participant was watching at any given time) was averaged over all participants. The predictor data of all three datasets were combined to get a stable clustering result over different types of movie stimulus. Correlation matrices were calculated for all intended analysis time scales (200 ms, 500 ms, 1000 ms, 2000 ms, and 4000 ms time windows) and the average correlation matrix over these time window matrices was generated to get stable clustering results over different temporal scales. The average correlation matrix was clustered hierarchically using the default UPGMA algorithm (https://www.rdocumentation.org/packages/stats/versions/3.6.2/topics/hclust). The suitable number of clusters was chosen using visual inspection of the clustered correlation matrix (**Figure SI-5**). The goal was to have a decent reduction in predictor correlations while preserving interpretability of the features. Finally, cluster predictors were generated by averaging over the standardized predictors within each cluster. This resulted in the final set of 16 predictors.

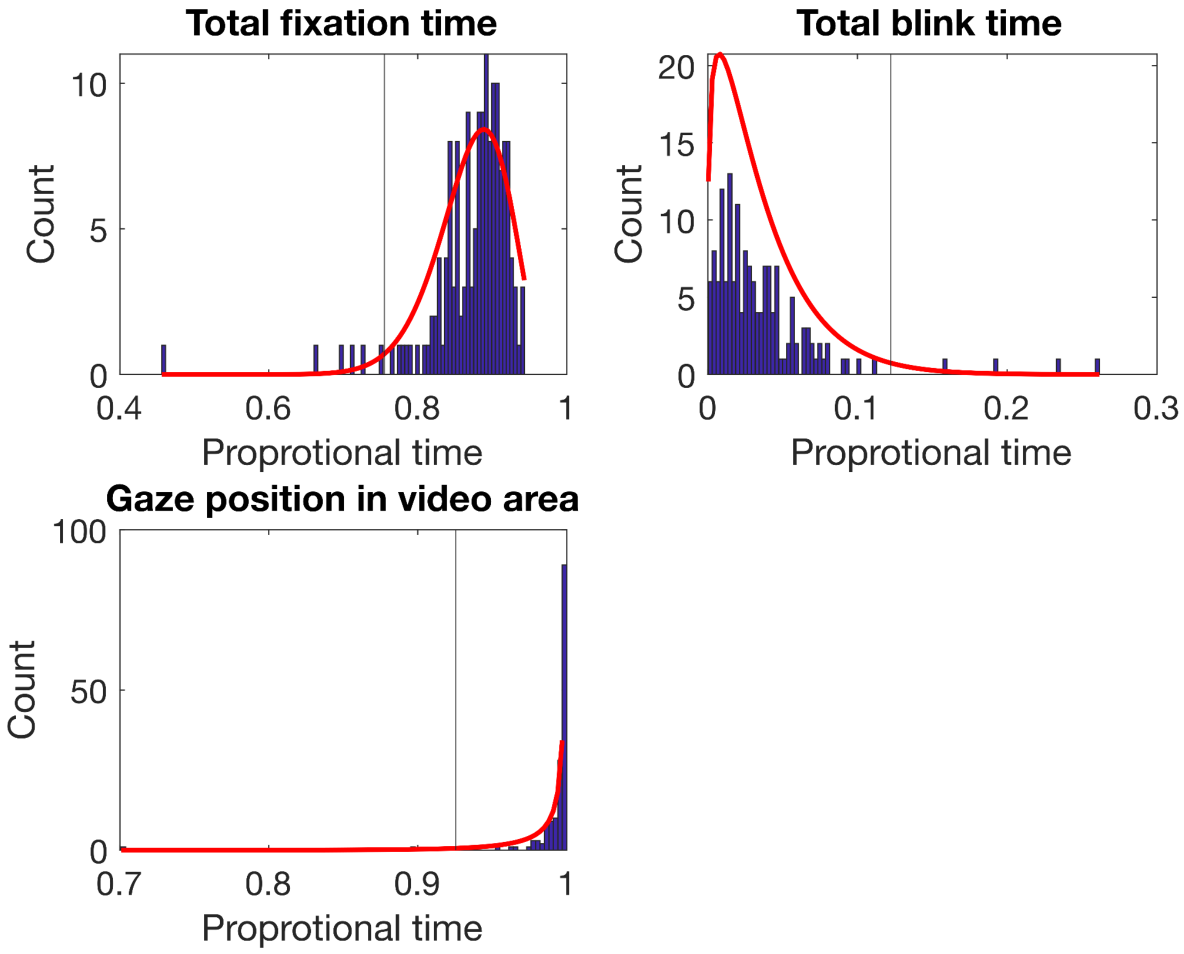


**Figure SI-6** Eye-tracking data quality control. The distributions visualize the total fixation and blink times (proportional to the total stimulus time) as well as the total time when the participant's gaze position was within the video area. Beta distributions (red lines) were fitted to the data and outlier participants were identified if their data fell into the 2% tail probability (vertical line) of the fitted distributions. Eight participants were excluded based on these data quality control.

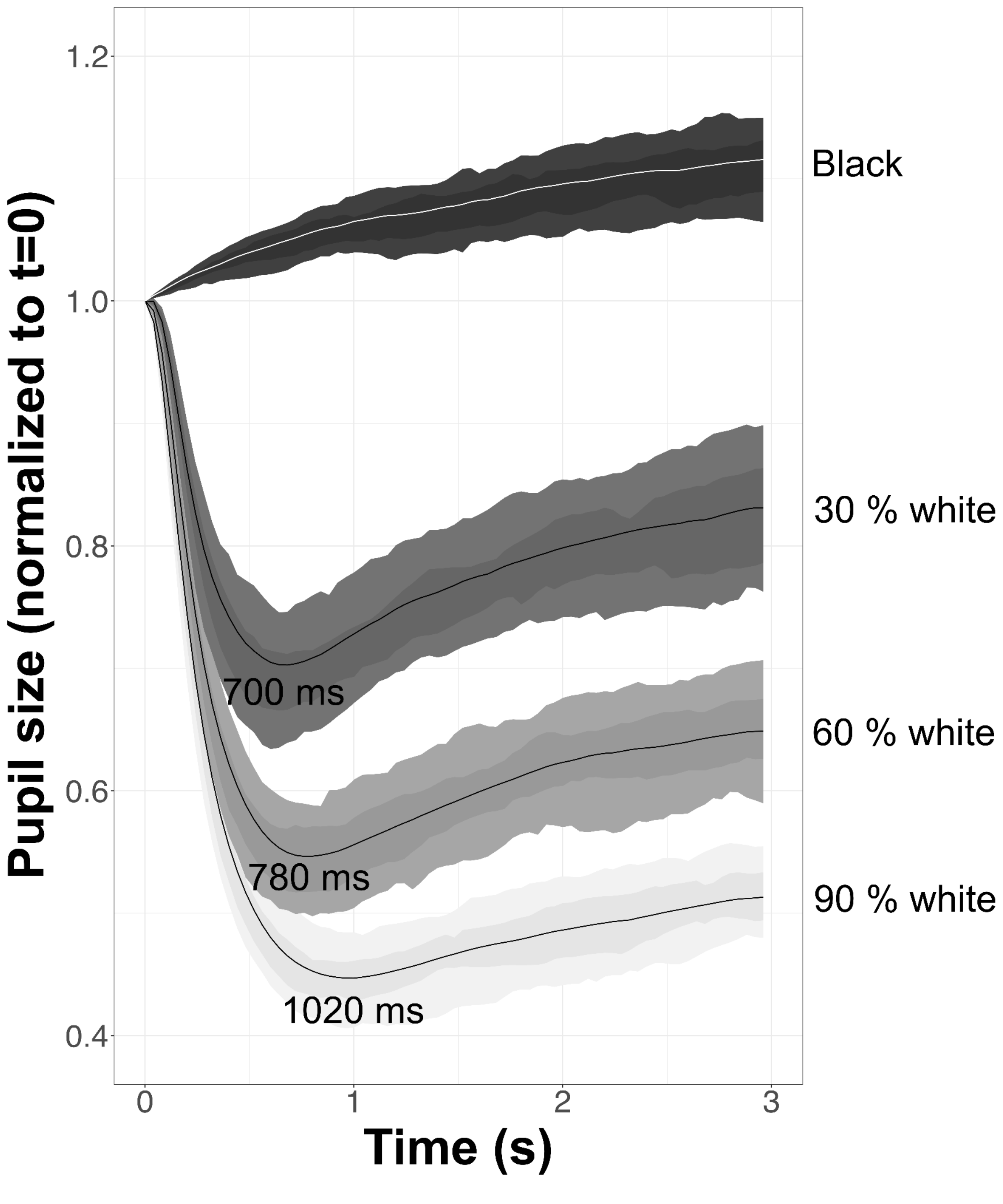


**Figure SI-7.** The pupillary light reflex of the participants for Experiment 1 participants. The line plots visualize the average pupillary responses for the luminance change of the presentation monitor. In the complementary pupillary light reflex experiment, the monitor screen was completely black at baseline and the screen luminance changed at time = 0. Pupil size decreased with increasing luminance and the response peaked around one second after the luminance change.

Random forest regression optimization

The tree size of the ensemble method and the number of branch splits in each tree was optimized within the training dataset using 80/20 train-test split. Correlation between the test split predictions and the actual values was used as the performance metric and computation time was also considered. In the Experiment 1 & 2 datasets the addition of trees drastically increased computation time with little increase in performance, while adding branches to the trees increased performance with a limited increase in computational time (**Table SI-2**). Since the optimization results were highly similar between the two datasets, the optimization was not repeated on the Experiment 3, where the optimization would have taken even more computing time. Based on these optimization results we chose to train random forest regression models with 50 trees and 63 branches in each tree.

| Experiment 1 | | | | Experiment 2 | | | |
|---|---|---|---|---|---|---|---|
| **Performance (test correlation)** | **Branches** | **Trees** | **Time (min)** | **Performance (test correlation)** | **Branches** | **Trees** | **Time (min)** |
| 0,5693 | 127 | 100 | 171,9 | 0,4348 | 127 | 50 | 602,3 |
| **0,5687** | **63** | **50** | **71,6** | 0,4338 | 127 | 100 | 1222,2 |
| 0,5657 | 127 | 50 | 87,4 | 0,4333 | 63 | 100 | 1051,7 |
| 0,5655 | 63 | 100 | 149,3 | **0,4332** | **63** | **50** | **539,5** |
| 0,5610 | 31 | 50 | 66,2 | 0,4320 | 31 | 100 | 911,1 |
| 0,5605 | 31 | 100 | 127,0 | 0,4317 | 31 | 50 | 448,4 |
| 0,5548 | 15 | 100 | 105,9 | 0,4312 | 15 | 100 | 758,6 |
| 0,5457 | 15 | 50 | 50,6 | 0,4280 | 15 | 50 | 378,0 |

**Table SI-2.** Random forest optimizations results. Performance was measured as the correlation between the predictions and true gaze probabilities in the test split of the training set. The results with the optimal parameters based on performance and computing time are indicated with boldface.
